# Egocentric boundary vector tuning of the retrosplenial cortex

**DOI:** 10.1101/702712

**Authors:** Andrew S. Alexander, Lucas C. Carstensen, James R. Hinman, Florian Raudies, G. William Chapman, Michael E. Hasselmo

## Abstract

The retrosplenial cortex is reciprocally connected with a majority of structures implicated in spatial cognition and damage to the region itself produces numerous spatial impairments. However, in many ways the retrosplenial cortex remains understudied. Here, we sought to characterize spatial correlates of neurons within the region during free exploration in two-dimensional environments. We report that a large percentage of retrosplenial cortex neurons have spatial receptive fields that are active when environmental boundaries are positioned at a specific orientation and distance relative to the animal itself. We demonstrate that this vector-based location signal is encoded in egocentric coordinates, localized to the dysgranular retrosplenial sub-region, independent of self-motion, and context invariant. Further, we identify a sub-population of neurons with this response property that are synchronized with the hippocampal theta oscillation. Accordingly, the current work identifies a robust egocentric spatial code in retrosplenial cortex that can facilitate spatial coordinate system transformations and support the anchoring, generation, and utilization of allocentric representations.

## Introduction

Spatial cognition is a critical component of intelligent behavior. The ability to effectively recall and navigate between known goals relies on stored representations of spatial interrelationships. Further, episodic experiences can be thought of as situated within a stored mental map indicating the places in which events occurred (Tulving, 1972). Spatial representations that support both navigation and episodic memory are observed in many brain regions, including the hippocampus (HPC) and medial entorhinal cortex (mEC), where neurons exhibit receptive fields that are correlated with the position or orientation of the animal relative to the array of locations and cues that define the structure of the outside world. This viewpoint-invariant coordinate system is commonly referred to as the allocentric reference frame (O’Keefe and Dostrovsky, 1971; Taube et al., 1990ab; Hafting et al., 2005).

Although it has been repeatedly shown that intact function of allocentric spatial circuits is critical for spatial memory and navigation (Morris et al., 1982; Taube et al., 1992; Steffenach et al., 2005), it is important to consider that all spatial information enters the brain via sensory organs and their corresponding processing streams. Accordingly, knowledge of the position of a prominent landmark and a neighboring goal location would be, at least initially, incorporated into a stored spatial map in egocentric coordinates relative to the animal itself (Andersen et al., 1983; Andersen et al., 1993; Andersen, 1997; McNamara and Rump, 2003; Burgess, 2006; Byrne et al., 2007; Bicanski and Burgess, 2018). Further, enacting navigational plans can be based upon stored allocentric representations but would ultimately require translation into sequences of actions anchored in an egocentric reference frame (e.g. one turns clockwise relative to their own previous orientation position; Byrne et al., 2007; Whitlock et al., 2008; Bicanski and Burgess, 2018).

Neural mechanisms by which egocentric and allocentric coordinate systems are interrelated are still the subject of intense examination. Computational models have predicted that cortical networks capable of integrating allocentric and egocentric information for either constructing or utilizing stored spatial representations require neurons with egocentric sensitivity to external locations (Byrne et al., 2007; Bicanski and Burgess, 2018). Most investigations into egocentric representations in unconstrained animals have focused on the neural substrates of path-integration, a navigational computation wherein self-location is approximated via continuous integration of angular and linear displacement (Mittelstaedt and Mittelstaedt, 1980; McNaughton et al., 2006). Neural correlates of these movement variables have been reported in several structures (McNaughton et al., 1994; Cho and Sharp, 2001; Whitlock et al., 2012; Kropff et al., 2015; Alexander and Nitz, 2015; Hinman et al., 2016; Wilber et al., 2017).

Only recently have externally-anchored egocentric representations that extend beyond self-motion been reported (Wilber et al., 2014; Peyrache et al., 2017; Wang, Chen, et al., 2018; Hinman et al., 2019, LaChance et al., 2019). Egocentric representations of this nature may anchor to environmental boundaries. Boundaries present a unique intersection between egocentric and allocentric coordinate systems as they have fixed positions that define the navigable allocentric space and simultaneously restrict the egocentric affordances of the agent such as what can be viewed or what motor plans can be executed. Importantly, environmental bounds or walls extend along large regions of an environment, and thus enable extended interaction from multiple allocentric or egocentric perspectives. Egocentric neural responses have now been reported in multiple areas such as lateral entorhinal cortex (Wang, Chen, et al., 2018), dorsal striatum (Hinman et al., 2019), and postrhinal cortices (LaChance et al., 2019). However, none of these regions possess the reciprocal interconnectivity between egocentric and allocentric spatial circuitry that might mediate bidirectional reference frame transformations.

From a connectivity standpoint, the retrosplenial cortex (RSC) is an excellent candidate to examine egocentric representations during navigation. Further, theoretical work has posited that RSC forms a computational hub for supporting coordinate transformations (Byrne et al., 2007; Clark et al., 2018; Rounds et al., 2018; Bicankski and Burgess, 2018). RSC is composed of two interconnected sub-regions, dysgranular (dRSC) and granular (gRSC), which have slightly different connectivity with cortical and subcortical regions (Shibata et al., 2009). dRSC (in mice agranular RSC) is positioned along the dorsal surface of the brain and possesses biased interconnectivity with association, sensory, and motor processing regions that code in egocentric coordinates (Vogt and Miller, 1983; van Groen and Wyss, 1992; Reep et al., 1994; Shibata et al., 2004; Wilber, Clark, et al., 2015; Yamawaki et al., 2016; Olsen et al., 2016; Hovde et al., 2019). In contrast, gRSC has strong reciprocal innervation with the hippocampal formation and associated structures that are primarily sensitive to the allocentric coordinate system (van Groen and Wyss, 1990; Wyss and van Groen, 1992; van Groen and Wyss, 2003; Miyashita and Rockland, 2007; Sugar et al., 2011; Kononenko and Witter, 2012; Czajkowski et al., 2013; Olsen et al., 2017; Yamawaki et al., 2019ab; Haugland et al., 2019).

Despite possessing dense reciprocal connectivity with numerous regions known to support spatial cognition, few reports have examined spatial response properties of neurons within the RSC. Most assessments of functional properties of RSC neurons have occurred in rodents performing track running tasks (Smith and Mizumori, 2012; Alexander and Nitz, 2015; Alexander and Nitz, 2017; Vedder et al., 2016; Mao et al., 2017; Mao et al., 2018; Miller et al., 2019). Track running experiments have revealed that RSC neurons exhibit spatial correlates with conjunctive sensitivity to allocentric and egocentric coordinate systems (among others) simultaneously (Alexander and Nitz, 2015). Conjunctive tuning of this type has been shown in modelling work to facilitate spatial coordinate transformations further supporting a role for RSC in the required transformation between these two spatial reference frames (Pouget and Sejnowksi, 1997; Bicanski and Burgess, 2018). However, grid cells, head direction cells, place cells, and other forms of well-characterized spatial receptive fields have primarily been examined in two-dimensional (2D) environments. Only a few experiments have studied RSC in similar conditions and all such reports have focused on head direction encoding (Chen et al., 1994ab; Cho and Sharp, 2001; Jacob et al., 2017).

To examine externally-referenced egocentric representations in RSC capable of supporting both navigation and reference frame transformations we recorded from both RSC sub-regions while rats freely explored familiar two-dimensional environments. We report that subsets of RSC neurons exhibit a variety of spatially-stable activation patterns in egocentric and allocentric coordinate systems. These findings support predictions from computational modeling related to translation between spatial reference frames and highlight important navigation related variables encoded in association cortex (Byrne et al., 2007; Bicanski and Burgess, 2018).

## Results

### RSC neurons exhibit stable spatial activity during free exploration

We recorded 555 neurons extracellularly in bilateral retrosplenial cortex (RSC) from male Long-Evans rats (n = 7) during free exploration. To enable comparisons between functional properties of neurons recorded in dysgranular (dRSC) versus granular (gRSC) sub-regions of RSC, we estimated tetrode placement and depth for each session (Figure 1a, n = 130 sessions, sFigure 1a-b). Of the total population, 41.5% (n = 230/555) were recorded from dRSC, 15.1% (n = 84/555) from the border between dRSC and gRSC, and 43.4% (n = 241/555) within gRSC. For baseline sessions, rats foraged for scattered reward in 1.25m^2^ square arenas with observable fixed distal cues.

**Figure 1.**
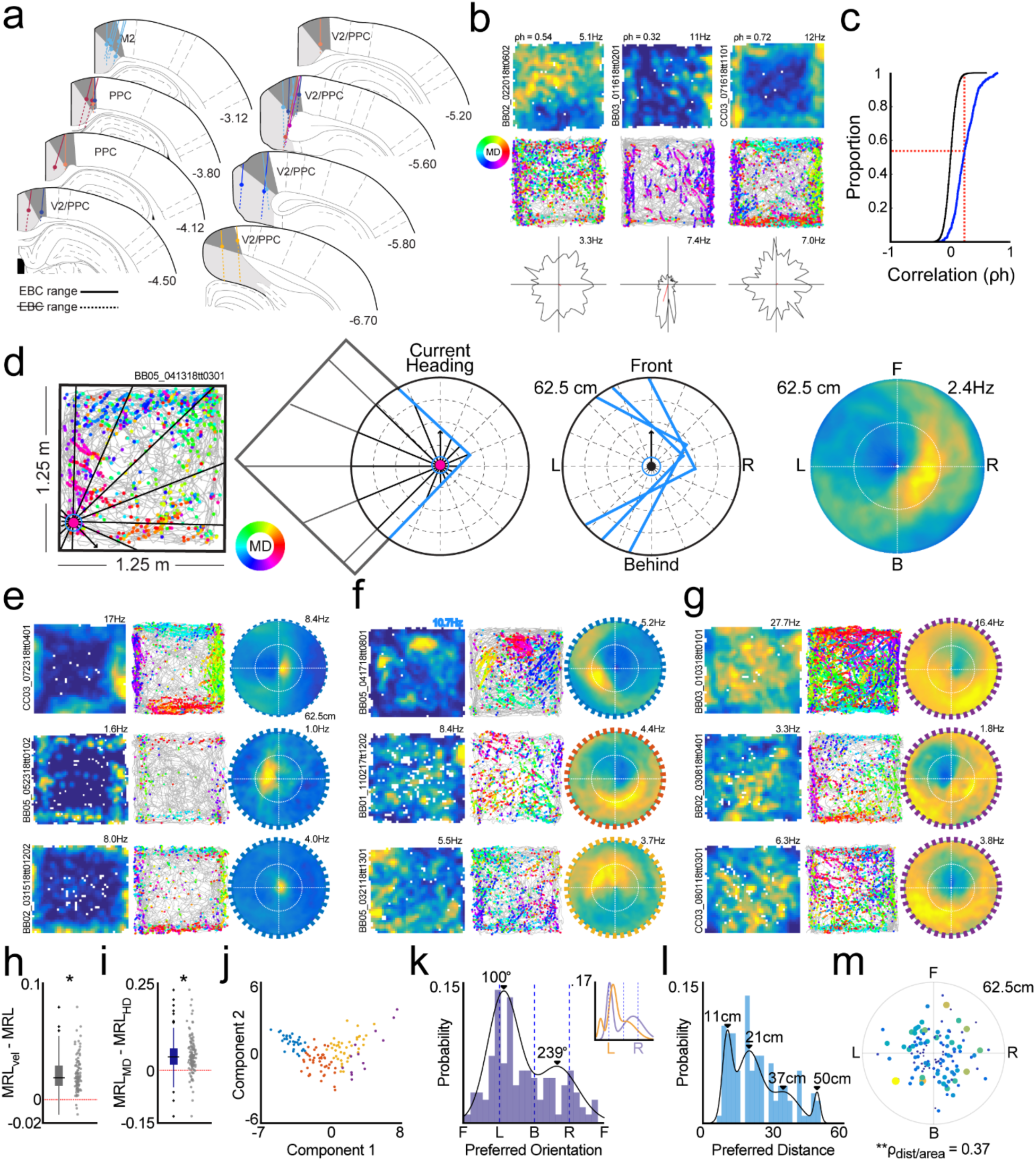
Egocentric boundary vector representations of RSC neurons during free exploration. **a**. Locations of RSC tetrode tracts where neurons with egocentric boundary sensitivity were observed. For each tetrode, solid lines indicate range where EBCs were recorded and filled circles indicate most ventral location of EBC observation. **b.** Example two-dimensional ratemaps (top), trajectory plots (middle), and head direction tuning plots (bottom) for three RSC neurons with significant stability in spatial firing. For trajectory plots, the position of the animal throughout the entire experimental session is depicted in gray. The location of individual spikes are shown with colored circles which indicate the corresponding movement direction of the animal according to the legend on the left. **c.** Cumulative density function depicts Spearman’s rho calculated after correlating 2D ratemaps taken from the first and second halves of each experimental session (blue). In black, distribution of spatial stability scores after randomly shifting spike trains relative to position. Red vertical line shows 99^th^ percentile of randomized distribution and its intersection with the real distribution of spatial stability. Percentage of neurons above red horizontal line have significant spatial stability. **d.** Schematic for construction of egocentric boundary ratemaps (EBRs). Left and middle panels, an example spike is mapped with respect to egocentric boundary locations in polar coordinates. Left, the movement direction of the animal is determined for each spike (vector with arrow) and the distance to wall intersections for all 360° are determined (sub-sample shown for clarity). Middle left, boundaries within 62.5cm are referenced to the current movement direction of the animal for a single spike. Middle right, example boundary positions for three spikes. Right, example egocentric boundary ratemap (EBR). **e.** Two-dimensional ratemaps, trajectory plots, and egocentric boundary ratemaps for 3 example RSC EBCs with animal-proximal receptive fields. **f.** Same as in e, but for 3 RSC EBCs with animal-distal receptive fields. **g.** Same as in **e** and **f**, but for 3 RSC EBCs with inverse receptive fields. **h.** Difference in strength (mean resultant length, MRL) of EBC tuning when a speed threshold was applied (MRL_vel_) versus no speed threshold (MRL). **i.** Difference in strength of EBC tuning when egocentric bearing was referenced to movement direction (MRL_MD_) rather than head direction (MRL_HD_). **j.** For neurons with significant egocentric boundary vector tuning, a scatter plot of first two principal component (PCA) scores calculated on multiple features of egocentric boundary ratemaps. Colors show 4 subsets of EBCs determined from K-means clustering on the same feature space as PCA and correspond to the colored boundaries around EBRs in **e-g**. **k.** Histogram of preferred orientation of receptive field across all RSC EBCs. Black line, probability density estimate from two-component Gaussian mixture model (GMM) on distribution of preferred orientation. Black triangles indicate peaks in GMM estimate. **l.** Same as **k**, but for preferred distance of all RSC EBC receptive fields. **m**. Polar scatter plot of preferred orientation versus preferred distance for the full RSC EBC population. Circle size indicates the area of the egocentric boundary vector receptive field.

RSC neurons exhibited complex firing rate fluctuations as rats randomly foraged within open arenas (Figure 1b). To assess the spatial stability of these representations for each neuron individually, we began by examining correlations between 2D spatial firing ratemaps constructed from first and second halves of each experimental session (Figure 1c). Across the full population of RSC neurons, 47.0% of cells (n = 261/555) had spatial correlations greater than the 99^th^ percentile (*ρ* = 0.23) of the distribution of correlations observed following 100 random shifts of the complete spike train for each neuron relative to spatial position (Figure 1c).

In some cases, RSC neurons with spatially anchored responses had slight differences in basic firing properties than those that were not spatially stable (sFigure 2b). Of particular interest and consistent with the presence of spatial receptive fields, RSC neurons with spatially reliable activity had significantly greater spatial coherence than non-stable cells (sFigure 2b-c). Spatially anchored firing patterns were also observed at more ventral recording sites where it was difficult to resolve whether the recording tetrode was in RSC or the cingulum bundle (sFigure 2d). Recordings from these sites were not included in the pool of RSC neurons for analysis.

**Figure 2.**
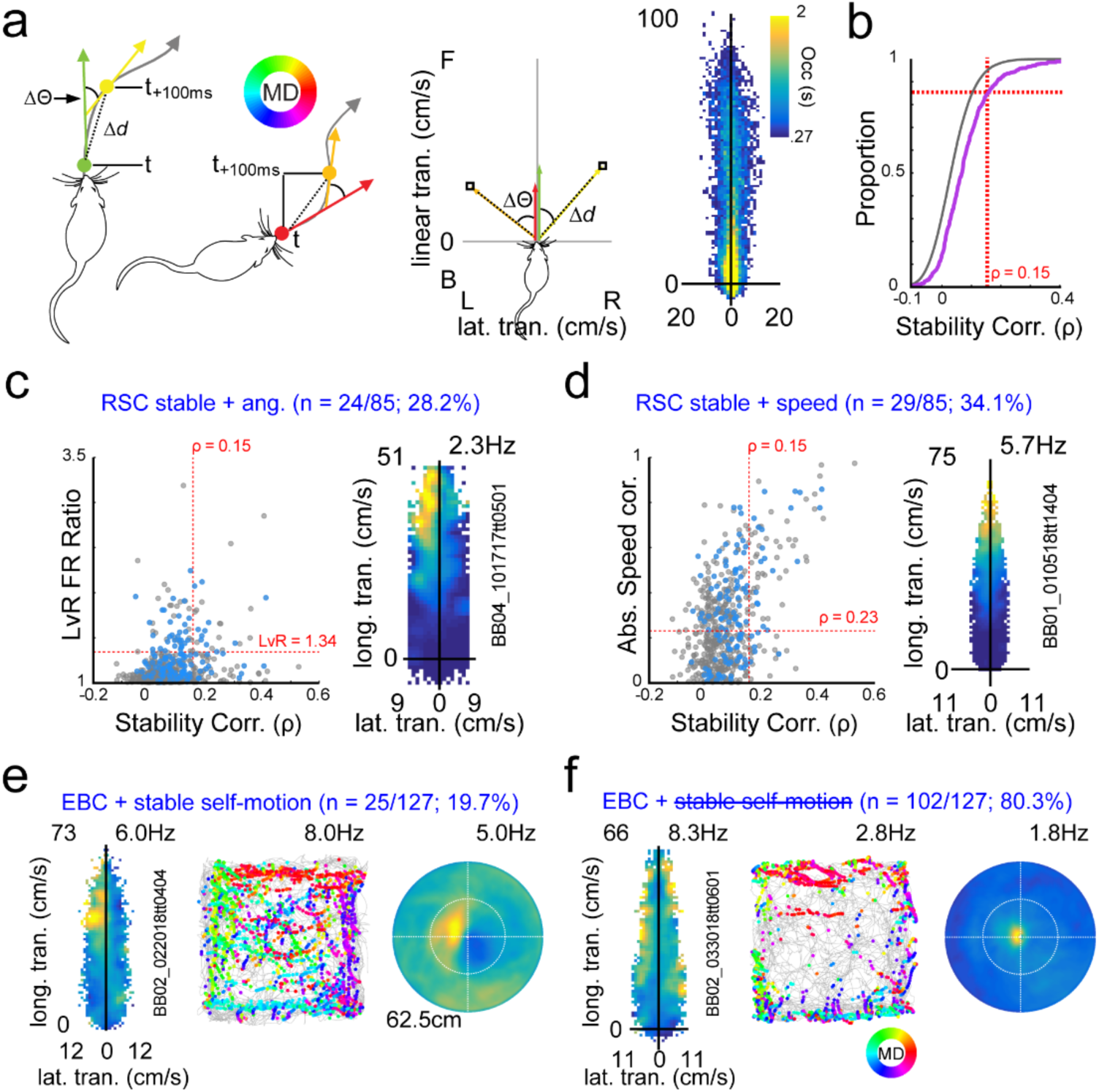
RSC egocentric boundary vector representations cannot be explained purely by self-motion correlates. **a.** Schematic of generation of self-motion referenced ratemaps. Left, example angular and distance displacements across 100ms temporal windows for two hypothetical position samples. Middle, corresponding lateral and longitudinal displacement for left examples in self-motion referenced coordinates. Right, heat map depicting mean occupancy in seconds for lateral and longitudinal displacement combinations across a complete experimental session. **b.** In pink, cumulative density functions for self-motion ratemap stability values (Spearman’s ρ) for all RSC neurons (randomization in gray). Red vertical line shows 95^th^ percentile of randomized distribution and its intersection with the real distribution of spatial stability. Percentage of neurons above red horizontal line have significant spatial stability. **c.** Left, spatial stability score (x-axis) versus absolute ratio of activity on left versus right halves of self-motion ratemaps (y-axis) for all RSC neurons. Blue dots correspond to identified RSC EBCs. Red lines and corresponding values correspond to 99^th^ percentiles of randomized distributions for both metrics. Neurons with values in upper right region were determined to have significant angular displacement tuning. Right, example RSC neuron with significant firing rate modulation for counterclockwise movements. **d.** On left, same as in **c**, but for spatial stability score versus absolute correlation between mean firing rate and animal speed. Right, example RSC neuron with significant firing rate modulation as a function of animal speed. **e.** Example RSC EBC with stable self-motion correlates**. f.** Example RSC EBC with non-stable self-motion correlates.

### Egocentric boundary vector responsivity of RSC

Of neurons with stable spatial firing in the open field, several had receptive fields that were qualitatively proximal to environmental boundaries (Figure 1b, **right**). Inspection of the relationship between each spike and the movement direction of the animal revealed that these responses manifested when the animal was oriented in a similar manner relative to any wall, suggesting that the receptive field was defined in an egocentric manner. As such, these responses were reminiscent of egocentric boundary cells (EBCs) recently reported in the dorsal striatum (dStr, Hinman et al., 2019), lateral entorhinal cortex (LEC, Wang, Chen, et al., 2018), and postrhinal cortex (POR, LeChance et al., 2019). To test this explicitly we constructed egocentric boundary ratemaps (EBRs) using procedures previously described (Hinman et al., 2019; Figure 1d). Briefly, for each behavioral frame, the distance to the nearest wall in each 3° offset from the animal’s movement direction was calculated (Figure 1d). The same process is repeated for the position of each spike from each neuron, and then ratemaps in polar coordinates were constructed by dividing the number of spikes by the total behavioral occupancy in seconds.

From each EBR, we computed the mean resultant length (MRL) of angular tuning as well as the absolute difference in angular tuning direction and distance between first and second halves of the baseline session. RSC cells were determined to exhibit significant egocentric boundary sensitivity if they met the following criteria: 1) they had a MRL for the first and second halves of the session that were greater than the 99^th^ percentile of the distribution of resultants computed following repeated shifted spike train randomizations, 2) had an absolute difference of mean directional tuning between halves of the baseline session that was less than 45 degrees, and 3) had an absolute difference in preferred distance tuning (described below) between halves that was less than 75% of the preferred distance tuning computed from the full baseline session.

Using these metrics, 15.0% (n = 83/555) of RSC neurons were determined to be EBCs (Figure 1e-g). When a speed threshold was applied (> 5 cm/s), a greater population of RSC neurons reached EBC criterion (22.9%, n = 127/555), which we utilize for further analyses. Overall, application of a speed threshold increased the mean resultant length for nearly all EBCs (Figure 1h, median MRL difference = 0.02, IQR = 0.01-0.03; Wilcoxon Signed Rank, z = 7.73, p = 1.1×10^−14^). This result suggested that the egocentric receptive field of EBC neurons is defined by the movement direction of the animal rather than head direction, which can be computed even when the animal is motionless. Indeed, when EBCs were assessed using head direction instead of movement direction the MRL significantly dropped (EBCs MRL with MD = 0.13, IQR = 0.10-0.18; EBCs MRL with HD = 0.10, IQR = 0.06-0.14, Wilcoxon sign rank test for zero difference, z = 8.04, p = 8.75×10^−16^).

### Properties of RSC egocentric boundary vector receptive fields

Sub-populations of RSC EBCs exhibited either increased or decreased activation when the animal occupied a particular orientation and distance relative to environmental boundaries (Figure 1e-g). In accordance with prior literature, we refer to those neurons that were inhibited as inverse EBCs (iEBCs, Figure 1g) and neurons with excitatory receptive fields as EBCs (Figure 1e-f). K-means clustering on numerous EBC features (see methods) yielded four qualitatively coherent groupings of EBC receptive field sub-types that were characterized post hoc: 1) 24.4% (n=31/127) were animal-proximal with a small receptive field (Figure 1e, **blue border**), 2) 33.9% (n=43/127) were animal-distal with a larger and/or potentially noisier receptive field (Figure 1f, **orange border**), 3) 31.5% (n = 40/127) were animal-distal with a larger receptive field (Figure 1f, **yellow border**), and 4) 10.2% (n = 13/127) had large inverted EBC receptive fields (Figure 1g, **purple border**). Principal component analysis (PCA) on this same feature space showed that the first two components accounted for 55.6% and 10.8% of the variability, respectively. A comparison of PCA scores for these two components across all RSC EBCs clustered using K-means did not yield distinct boundaries between sub-populations, instead revealing a continuum of EBC receptive fields (Figure 1j).

Identification of the center of mass of EBC receptive fields revealed a bimodal distribution of preferred orientations that was best fit by a two component Gaussian mixture model (GMM) with means of 100° (L) and 239° (R) relative to directly in front of the animal (Figure 1k; Number of GMM components determined via minimizing AIC). Although EBCs were recorded in both hemispheres there was no obvious relationship between the preferred orientation and the hemisphere in which the neuron was recorded, indicating a lack of lateralization of EBC response properties (Figure 1k, inset). The distribution of preferred distances was best described by a four component GMM with means of 11cm, 21cm, 37cm, and 50cm centimeters (Figure 1l). The size of EBC receptive fields increased as a function of the preferred distance of the egocentric vector indicating that the resolution of the representation was dependent on proximity to boundaries (Figure 1m, **Spearman’s *ρ* = 0.37, p = 2.57×10^−5^**). The presence of EBCs with preferred distances distal to the animal suggested that the EBC response property was neither dependent upon physical interaction with arena borders nor could be purely posture related which is known to modulate cortical neurons (Mimica, Dunn, et al., 2019).

### EBC responses are localized within dysgranular RSC but lack topographic organization

Egocentric boundary vector sensitivity was primarily observed in dRSC, where 37.0% (n = 85/230) of neurons recorded were classified as EBCs (sFigure 1c). In contrast, EBCs were observed in 10.0% (n = 24/241) of gRSC and 21.4% (n =18/84) of intermediary area cells between the two sub-regions (dgRSC, see methods and sFigure 1a). By and large, the distribution of EBCs amongst RSC sub-regions was consistent across animals (sFigure 1c). The EBC response property was observed across a wide range of A/P coordinates spanning a majority of RSC but possessed no further anatomical organization beyond sub-region specificity (range = 2.9 - 6.8mm relative to bregma, sFigure 1d). The distribution of spike waveform widths across all RSC neurons was bimodal with identified EBCs primarily found in the cluster of neurons with longer duration waveforms (sFigure 1e, K-means clustering on waveform width, cluster 1 median = 0.18s, IQR = 0.15 - 0.23s, EBCs in cluster 1, n = 15/127, 11.8%; cluster 2 median = 0.31s, IQR = 0.21 - 0.34s; EBCs in cluster 2, n = 112/127, 88.2%). Further, EBCs had overall low mean firing rates (sFigure 1e, EBCs = 1.68Hz, IQR = 0.96 - 2.72Hz, not-EBCS = 3.55Hz, IQR = 1.15 - 8.47Hz). Taken together, the EBC sub-population was determined to be primarily composed of putative principal neurons suggesting that the EBC signal is propagated across RSC sub-regions or into other brain regions.

EBCs could often be simultaneously recorded which enabled an analysis of potential topography in the distribution of preferred distance and orientation of the egocentric boundary vector. Overall, 129 pairs of RSC EBCs were co-recorded across 31 sessions. Of these pairs, 30.2% (n = 39/129) were recorded on the same tetrode (sFigure 3a), while the remaining 69.8% of EBC pairs (n = 90/129) were concurrently recorded on different tetrodes (sFigure 3b).

**Figure 3.**
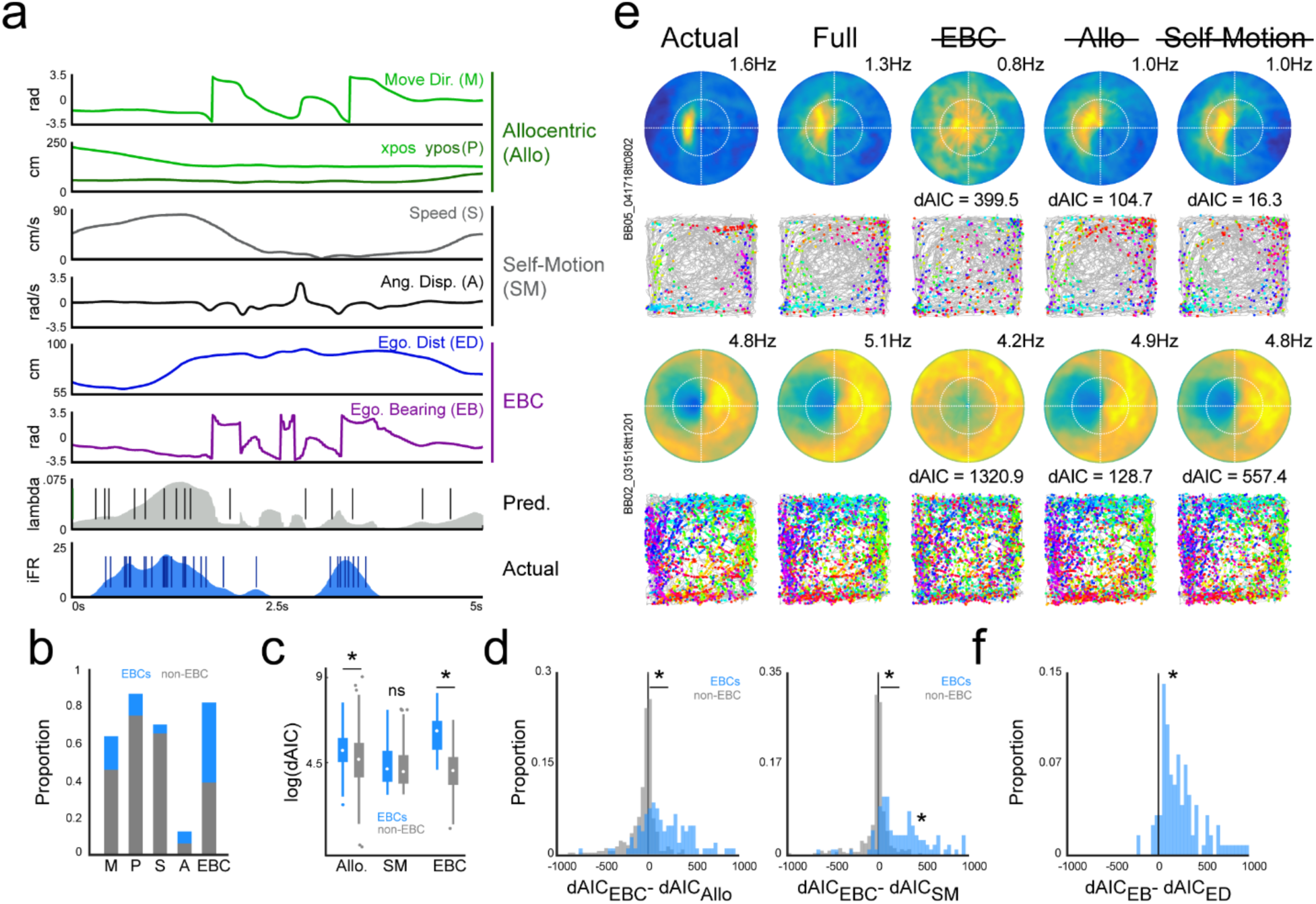
Egocentric vector tuning is more robust than allocentric or self-motion correlates using a generalized linear modelling framework. **a.** Example GLM predictors composing allocentric, self-motion, and egocentric vector classes with corresponding actual and predicted firing rates and spike trains over a five second window. **b.** Proportion of non-EBC RSC neurons (gray) and EBC RSC neurons (blue) exhibiting significant sensitivity to each predictor individually. M, movement direction; P, x- and y- position; S, speed; A, angular displacement; EBC, egocentric boundary. **c.** Boxplots depicting median and quartiles of log-transformed difference of Akaike information criteria scores (dAIC) for models with all allocentric, self-motion, or egocentric vector predictors removed (blue bars, EBCs; gray bars, non-EBCs). Larger dAICs indicate greater error in model fit with removal of a predictor class. **d.** Comparison of dAIC scores for models with egocentric vector versus allocentric predictors removed (left) or egocentric vector versus self-motion predictors removed (right) for EBCs (blue) and non-EBCs (gray). Rightward shifts indicate greater error in model fit for models with removed egocentric vector predictors. **e.** For two example RSC EBCs, predicted GLM spike trains from all models were utilized to construct egocentric boundary ratemaps and trajectory plots. Left column, actual egocentric boundary ratemap and corresponding trajectory plot. Second column, for the same cell, an egocentric boundary ratemap and corresponding trajectory plot for the generalized linear model constructed using all egocentric vector, allocentric, and self-motion predictors. Final three columns, egocentric boundary ratemaps and trajectory plots for each reduced model and corresponding dAIC scores. **f.** Comparison of dAIC scores for models with the egocentric bearing versus the egocentric distance removed reveal greater impact of egocentric bearing for EBCs.

To assess whether there was organization to preferred orientation and distance as a function of proximity of two EBCs (i.e. observed on same or different tetrodes), we next calculated the difference in receptive field center of mass for both angular and distance components for all pairs. Although the preferred distance of EBCs on the same tetrode was trending towards greater similarity (sFigure 3c), neither preferred orientation nor distance was statistically different for EBCs recorded on the same versus different tetrodes (sFigure 3c-3d, absolute difference in preferred distance (PD) same tetrode = 7.5cm, IQR = 5 - 21.88cm; absolute difference in PD different tetrode = 12.5cm, IQR = 5 - 22.5cm; Wilcoxon rank sum test, z = - 1.16, *p* = 0.24; difference in preferred orientation (PO) same tetrode = 33°, IQR = -43.5 – 78.75°; difference in preferred orientation (PO) different tetrode = 34.5°, IQR = -6 – 111°; Wilcoxon rank sum test, z = -0.04, *p* = 0.97). Accordingly, we conclude that there is a lack of topographic organization of egocentric boundary vector tuning in the RSC.

### Egocentric boundary vector tuning in secondary motor cortex and posterior parietal cortex but not medial entorhinal cortex

In 3 animals, a subset of more anterior recording tetrodes were positioned in secondary motor cortex (M2, from bregma: AP: -1.1 to -2.9mm, ML: ±0.8 to 1.2mm) and 56 neurons were recorded there (sFigure 4a). Of M2 neurons, 21.4% reached EBC criterion (n = 12/56, sFigure 4b). Similarly, 95 neurons across 5 rats were recorded in posterior parietal cortex (PPC or V2, from bregma: AP: -3.7 to -5.9mm, ML: ±1.5 to 2.4mm) and a sub-population of 9.5% (n = 9/95) reached EBC criterion (sFigure 4c-d). EBCs and iEBCs were observed in both structures and receptive fields had similar angular and distance distributions as those observed in RSC (sFigure 4e-f). In contrast, only 2.4% (n = 7/297) of medial entorhinal cortex (mEC) neurons recorded in similar conditions reached EBC criterion, indicating that the egocentric vector signal was generally not present within the region (sFigure 4g, Wang, Chen, et al., 2018).

**Figure 4.**
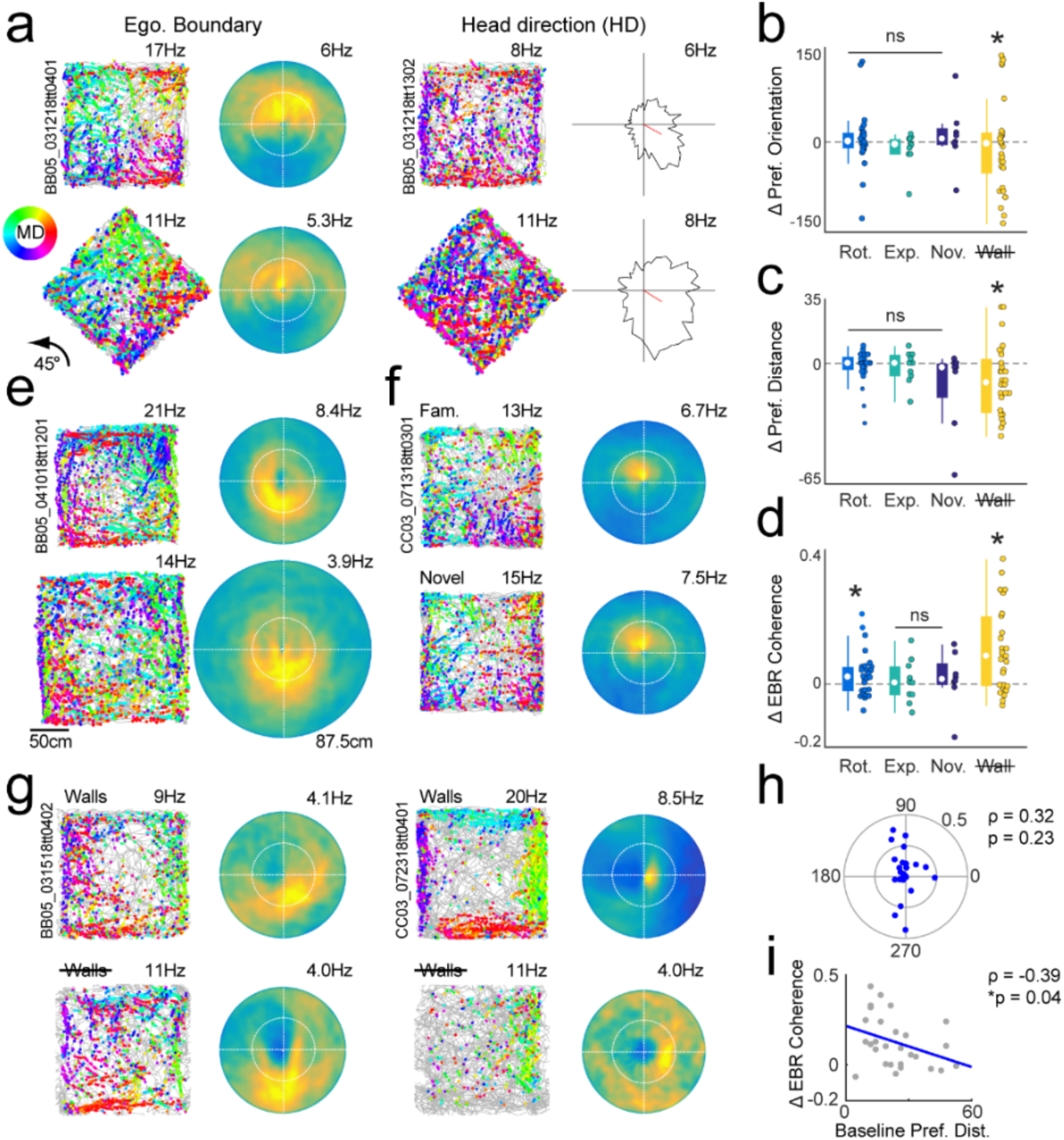
EBCs are anchored to local boundaries, respond in novel environments, and lose sensitivity in arenas without explicit borders. **a.** Left, trajectory plot and egocentric boundary ratemap (EBR) for an example EBC with similar egocentric boundary vector tuning in baseline experimental session (top) and a second session in an environment rotated 45° (bottom). Right, example head direction neuron sustains directional tuning across both conditions. **b.** Preferred orientation of EBC receptive fields in all arena manipulation sessions subtracted from preferred orientation in baseline sessions. **c.** Preferred distance of EBC receptive fields in all arena manipulation sessions subtracted from preferred distance in baseline sessions. **d.** EBC receptive field coherence in all arena manipulation sessions subtracted from receptive field coherence in baseline sessions. **e.** Trajectory plot and EBR for an example EBC with similar egocentric boundary vector tuning in baseline experimental session (top) and a second session in an expanded arena (bottom). **f.** Trajectory plot and EBR for an example EBC with similar egocentric boundary vector tuning in baseline experimental session (top) and a second session in a novel arena (bottom). **g.** Trajectory plot and EBR for two example EBCs between baseline session (top) and session with walls removed (bottom). Left EBC has a more distal receptive field and exhibits similar egocentric boundary vector tuning. Right EBC has a more proximal receptive field and has disrupted tuning in arena with no walls. **h.** For EBCs recorded in arenas without walls, the preferred orientation at baseline plotted against the change in EBC receptive field coherence between the two sessions. **i.** Same as **h,** but for change in coherence as a function of baseline preferred distance.

### EBC responsivity is not explained by self-motion correlates

In free exploration, spatial locations near environment boundaries uniquely restrict the behavioral affordances of the animal. Many observed EBC receptive fields were proximal to the rat, firing only when the animal was close to boundaries and thus most limited in its possible actions. We next tested whether the manifestation of egocentrically referenced boundary vector tuning was in actuality reflective of self-motion related firing that was stereotyped near borders.

We began by constructing self-motion referenced firing rate maps during open field sessions (Chen et al., 1994; Whitlock et al., 2012). The angular difference between movement direction (Δθ) and the Euclidian distance in two-dimensional location (Δd) was calculated across a sliding 100ms window for every position of the animal throughout a free exploration session (Figure 2a, **left**). These displacement values were converted to Cartesian coordinates referenced to the previous location of the animal at each step, thus producing a map of the distance and direction of movement of the animal for all position samples within the exploration session (Figure 2a, **middle and right**).

Firing rate as a function of these displacement values are presented for example RSC neurons in Figure 2c-f. The zero-line intersection indicates the position of the animal at the beginning of each 100ms window and the x and y-axes reflect displacement in lateral and longitudinal dimensions, respectively. Thus, values to the right of the vertical zero line reflect the activity of the neuron when the animal moved to the right relative to the previous position and direction of its body axis and the distance that the action took the animal is reflected in the position of the value along the y-axis.

To quantify the stability of self-motion tuning, we correlated self-motion ratemaps for each neuron that were individually computed from interleaved temporal epochs (1s in duration) within the free exploration session. 15.3% (n = 85/555) of RSC neurons exhibited self-motion related activity that had greater stability than the 95^th^ percentile of the distribution of stability correlation values calculated following permutation tests (Figure 2b). Of this sub-population, 28.2% (n = 24/85) had firing rate modulation that was biased for leftward or rightward movements (Figure 2c), while 34.1% (n = 29/85) were sensitive to longitudinal movements consistent with speed tuning (Figure 2d). Of the EBC population, 19.7% (n = 25/127) met the stability criteria, indicating that a small sub-population of neurons exhibiting egocentric boundary vector tuning had stable self-motion correlates (Figure 2e). However, the vast majority of RSC EBCs did not exhibit self-motion correlates confirming that egocentric boundary vector tuning was primarily not an epiphenomenon of movement related activity near borders (Figure 2f).

Beyond EBCs, the present analysis demonstrated overall limited self-motion tuning in RSC during free exploration. This observation shines new light on previously reported turn-sensitive neurons in RSC during track running tasks (Alexander and Nitz, 2015). In prior work, the magnitude of clockwise or counterclockwise activation during track running was demonstrated to be generally insensitive to the magnitude of angular velocity on a trial-by-trial basis. In combination with the lack of self-motion tuning during free foraging observed here, the results collectively suggest that reported egocentric correlates in RSC are externally-referenced and unrelated to the speed of angular movement.

### Generalized linear models demonstrate robust egocentric vector tuning of RSC EBCs

Self-motion is necessarily conflated with egocentric boundary vector tuning because the response primarily manifested during movement (Figure 1h-i). An EBC may exhibit stable firing rate fluctuations as a function of self-motion that are driven by the egocentric boundary vector receptive field, not the action state of the animal. For example, an EBC with a receptive field to the animal’s left may also show self-motion tuning for clockwise movements as a result of the animal being more likely to turn clockwise when there is a wall occupying the egocentric receptive field. Yet, the same neuron may not be activated when the animal turns clockwise in other locations within the environment that do not satisfy the egocentric boundary vector. Thus, although initially informative, a different approach was required to tease out the influence of self-motion and other potential spatial covariates on EBC activity patterns.

We next implemented a generalized linear model (GLM) framework to predict the probability of spiking at each time point as a function of the relative influence of multiple allocentric, self-motion, or EBC-related predictors (Figure 3a). Allocentric predictors included the movement direction of the animal and x- and y-position within the arena. Self-motion related predictors included linear speed and angular displacement (i.e. the differential of animal movement direction in 100ms windows).

EBC-related predictors were more complicated as a single position sample or spike possessed relationships to multiple locations along boundaries simultaneously. Accordingly, the EBC predictor could take many forms. To minimize the number of sub-predictors, EBC predictors were composed of the animal’s distance and egocentric bearing to the center of the arena. Unlike arena boundaries, the center of the arena is a single coordinate that can be described as a function of individual angular and distance components or their conjunction for each position sample (sFigure 5a). Critically, EBCs were found to exhibit robust egocentric bearing and distance tuning to the center of the arena making the predictor a reasonable counterpart to referencing single unit activity to arena walls (sFigure 5c-d).

**Figure 5.**
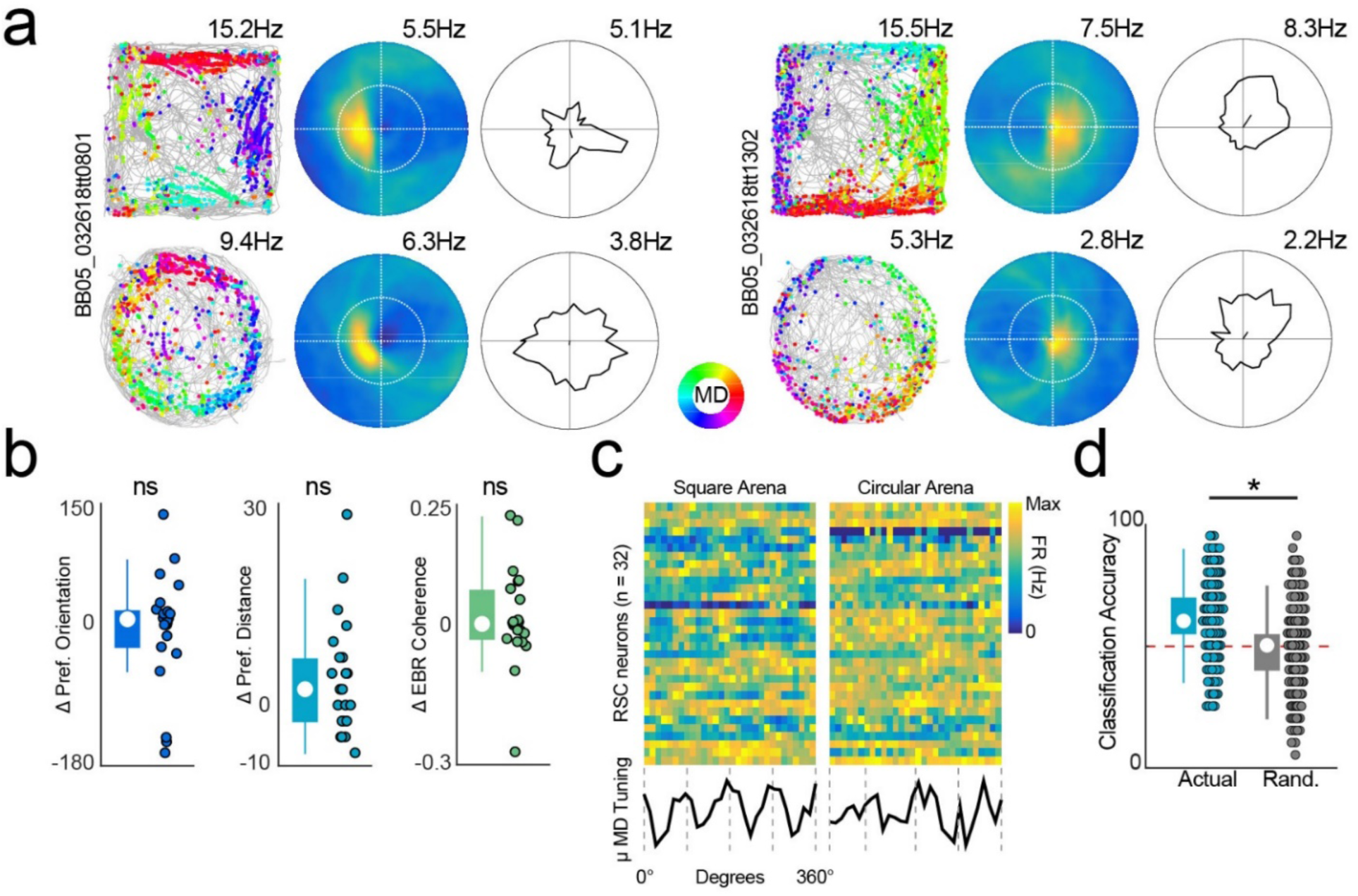
RSC EBCs are insensitive to environmental geometry which generates a directional representation of environment shape. **a.** Trajectory plots, egocentric boundary ratemaps (EBR), and movement direction tuning plots for two example RSC EBCs for experimental sessions in a square (top) and circular environment (bottom). **b.** Preferred orientation, preferred distance, and EBC receptive field coherence from recording sessions in the circular arena subtracted from the corresponding metrics in baseline sessions. **c.** Movement direction tuning plots for all RSC neurons in the square arena (left) and circular arena (right). Color depicts intensity of activation (blue is zero firing rate, yellow is maximum firing rate). Bottom in black, the average movement direction tuning across the full population of RSC neurons for the square and circular environments. Gray dashed lines depict 90° axes. **d.** Arena classification accuracy for linear discriminant analysis on movement direction tuning from **c**. In teal, actual classification. In gray, classification after randomizing arena identity. Red dashed line is statistical chance.

To test the impact of each predictor individually, we first generated a complete model fit using the full complement of predictors. Next, we dropped each covariate individually and assessed the decrement to model fit using the difference in Akaike information criterion and corresponding log-likelihood ratio tests (AIC, see methods). Figure 3b depicts the proportion of RSC neurons that had significant sensitivity to each individual predictor, split into the sub-populations of cells that were or were not classified as egocentric boundary sensitive using strength of bearing tuning and reliability as above and in previous work (Hinman et al., 2019). GLM output corresponded with the initial identification of EBCs using these metrics. Of EBCs, 81.9% (n = 104/127) had significant decrements to model fit when conjunctive EBC predictors were removed from the model (Figure 3b).

Using the GLM, 70% (n = 89/127) of EBCs were significantly modulated by linear speed, more than doubling the percentage of neurons that were identified as velocity sensitive from self-motion ratemap analyses that utilized a 100ms temporal window (Figure 2d). GLM analyses yielded only 12.6% (n = 16/127) of RSC EBCs sensitive to angular displacement, thus further confirming that egocentric boundary vector sensitivity was not epiphenomenal to turning actions near arena bounds.

Although neurons previously characterized as possessing egocentric boundary vector tuning were accurately detected with the GLM, numerous other predictors were shown to also significantly co-vary with spiking activity (Figure 3b). A number of neurons that were initially characterized as non-EBCs had significant model decrements when the EBC-related predictors were excluded (38.8%, n = 141/363 after removing non-EBC interneurons that had mean firing rates > 15Hz; Figure 3b). However, as the GLM will zero-weight a non-contributing predictor, the removal of a single covariate can only serve to decrement model fit while adding a predictor can only improve model fit. Accordingly, a significant log-likelihood test following removal of a predictor does not fully demonstrate the relative impact of that predictor in the full model.

We next assessed the overall influence of each predictor class (allocentric, self-motion, and EBC-related) on model fit by constructing a nested GLM, dropping each predictor class, and then making comparisons between resulting model fits (Kraus et al., 2013). Figure 3c depicts the difference in model fit (dAIC) for both EBCs and non-EBCs between the full model and reduced models with all allocentric, self-motion, or egocentric boundary predictors removed. Larger dAIC values indicate greater impact of the predictor class within the full model. Models without all allocentric or EBC predictors had significant differences in fit between EBCs and non-EBCs (Figure 3c, Kruskal-Wallis, χ^2^ = 275.67, p = 1.69×10^−57^, post-hoc Scheffe tests, p<0.0001). There was no difference between these sub-populations of RSC neurons for the removal of self-motion covariates from the model further supporting that EBCs were not more sensitive to speed or angular displacement than the remainder of the RSC population (Figure 3c, Kruskal-Wallis w/ post-hoc Scheffe tests, p=0.95).

A clear divergence emerged in the importance of EBC-related predictors for the EBC and non-EBC sub-populations. As reflected in the difference in magnitude of dAIC, EBC predictors had greater impact than either allocentric or self-motion predictors for the EBC population (Figure 3d, **blue;** dAIC_EBC_ - dAIC_Allo_ for EBCs = 203.5, IQR = 21 – 475.8; dAIC_EBC_-dAIC_SM_ for EBCs = 259.2, IQR = 65.6 – 541.1) than for the non-EBC population which had similar dAIC scores (near 0) for models lacking EBC predictors and other predictor classes (Figure 3d, **gray;** dAIC_EBC_ - dAIC_Allo_ for non-EBCs = -14.9, IQR = -110.4 – 10.2; dAIC_EBC_-dAIC_SM_ for non-EBCs = - 1.33, IQR = -29.6 – 26.4). Overall, the impact of EBC-related predictors relative to other predictor classes was significantly greater for EBC versus non-EBC sub-populations (dAIC_EBC_ - dAIC_Allo_ for EBCs versus non-EBCs, Wilcoxon rank sum, z = -10.8, p = 2.51×10^−27^; dAIC_EBC_- dAIC_SM_ for EBCs versus non-EBCs, Wilcoxon rank sum, z = -12.3, p = 9.33×10^−35^).

This result suggested that although models without allocentric or self-motion predictors could yield significantly decreased model fit, the vast majority of EBC neurons were significantly more impacted by EBC predictors. Two example EBCs in Figure 3e illustrate this point, wherein a spike train generated from the output of each model was used to construct trajectory plots and egocentric boundary ratemaps. In both cases, the model lacking egocentric orientation and distance information yields a trajectory plot and egocentric boundary ratemap that is substantially poorer at reconstructing the actual data than any other reduced model.

Although egocentric predictors were the dominant influence on EBC activation, nearly all EBCs (98.4%, n = 125/127) were statistically impacted by the removal of more than one predictor category. In this manner the GLM analyses revealed that RSC EBCs were conjunctively sensitive to the position of arena boundaries in egocentric coordinates and allocentric heading or location simultaneously. This feature of EBC responsivity is consistent with theoretical work proposing a transformation between egocentric and allocentric spatial representations within RSC (Bicanski and Burgess, 2018).

### GLM confirms vectorial representation

Use of the GLM framework provided an opportunity to verify that RSC neurons with egocentric boundary sensitivity actually formed vector representations of the relationships between environmental boundaries and the animal. By dropping out egocentric bearing and egocentric distance from the model individually, we were able to investigate the relative influence of the individual components of the egocentric boundary vector in isolation for each neuron.

Significant model decrements were observed in 100% (n = 127/127) of EBCs following removal of the egocentric bearing component and 74.8% (95/127) of EBCs were impacted by the removal of egocentric distance predictors. Overall, the magnitude of error to model fit was substantially greater when egocentric bearing was removed indicating that, although both distance and orientation components are critical for egocentric boundary vector responsiveness, the directional component more robustly drives neurons exhibiting this tuning preference (Figure 3f, difference in dAIC for egocentric bearing versus egocentric distance = 217.2, IQR = 77.5 – 440.4; Wilcoxon signed rank test, z = 9.3, p =1.3×10^−20^).

### EBCs respond to local not distal environmental features

Characterization of EBC properties and self-motion correlates were conducted in baseline sessions in which the open arena remained in a fixed location relative to the experimental room and fixed distal cues therein. We next conducted a series of experimental manipulations of the relationship between the familiar arena and the testing room in order to confirm that EBC response properties were defined by the relationships between environmental boundaries and the animal itself.

First, we rotated the open field 45 degrees to maximally disrupt correspondence between arena walls and distal walls or cues present within the recording environment to verify that EBC responses were anchored to local boundaries and not the broader recording room. Under these conditions we recorded a total of 65 RSC neurons (across 4 rats and 14 sessions) of which 40% (n = 26/65) had EBC sensitivity. Consistent with EBC responses being referenced to the rat, receptive fields in rotated arenas maintained the same orientation and distance with respect to the animal, even though arena boundaries now fell along completely different allocentric axes (Figure 4a-c; difference between baseline and rotated preferred orientation = 1.5°, IQR = -12 - 18°; Wilcoxon sign rank test, z = 0.53, p = 0.59; difference between baseline and rotated preferred distance = 0cm, IQR = -3.75cm – 3.75cm; Wilcoxon sign rank test, z = -0.26, p = 0.79). Although vector tuning remained intact, there were slight but significant changes to ratemap coherence between baseline and rotation sessions which suggested that the quality of the egocentric boundary receptive field was decremented across conditions (Figure 4d; difference between baseline and rotated ratemap coherence = 0.03, IQR = -0.02 – 0.06; Wilcoxon sign rank test, z = 2.27, p = 0.02).

Consistency in tuning could emerge if the allocentric map anchored to local boundaries rather than distal cues. This was not the case as a population of simultaneously recorded head direction cells (HD, n = 4; Figure 4a, **right**) exhibited consistent mean tuning across the rotated and non-rotated conditions (n = median tuning difference = -6.6°, maximum tuning difference - 12.4°). Accordingly, arena rotation experiments dissociated the directional component of EBCs from the allocentric reference frame of HD cells.

### EBC responsivity is anchored to boundaries not the center of the environment

RSC EBCs exhibited egocentric vector sensitivity to both arena boundaries and the center of the environment which we utilized to our advantage in GLM analyses (sFigure 5). This occurs because arena boundaries have a fixed relationship relative to the center of the environment. Accordingly, an obvious question is whether the egocentric boundary response is in actuality defined as an egocentric vector to the center of the arena. We addressed this possibility by comparing preferred orientation and distance for 10 RSC EBCs (of 33 total RSC neurons from 4 rats across 11 sessions) between baseline arenas and open fields expanded up to 1.75m^2^ (Figure 4e).

If EBC responses were anchored to boundaries, we anticipated that the orientation and preferred distance would remain consistent across both conditions. Conversely, if the receptive field was defined by a vector to the center of the arena then the distance component of the egocentric boundary vector would remain fixed to this point. In this scenario, the preferred distance would either move away from the animal in expanded arenas or potentially scale with the arena expansion. We observed that the preferred orientation, preferred distance, and ratemap coherence were not altered between baseline and expanded field sessions confirming that EBCs were indeed anchored to boundaries and not the center of the arena (Figure 4b-d, difference between baseline and expanded preferred orientation = -4.5°, IQR = -24 - 6°; Wilcoxon sign rank test, signed rank = 19, p = 0.46; difference between baseline and expanded preferred distance = 0cm, IQR = -7.5 – 5cm; Wilcoxon sign rank test, signed rank = 13.5, p = 0.56; difference between baseline and rotated ratemap coherence = 0.01, IQR = -0.03 – 0.07; Wilcoxon sign rank test, signed rank = 31, p = 0.77).

### EBC responsivity is stable in novel environments

Neurons within the broader neural spatial circuitry such as grid cells, head direction cells, and place cells, exhibit consistent, albeit remapped, spatial receptive fields in novel environments. We next questioned whether egocentric boundary vector tuned neurons of RSC would exhibit similar stability in their selectivity. We recorded 17 RSC cells including 8 EBCs in familiar then novel environment sessions (Figure 4f, 4 rats across 5 sessions). Neither distance nor orientation components of the egocentric boundary vector were altered in the novel environment relative to baseline illustrating that EBCs are not experience dependent and do not remap between environments (Figure 4b-c, difference between baseline and novel preferred orientation = 6°, IQR = -6 - 27°; Wilcoxon sign rank test, signed rank = 19, p = 0.47; difference between baseline and novel preferred distance = -2cm, IQR = -20 – 0cm; Wilcoxon sign rank test, signed rank = 2, p = 0.13). Coherence of EBC receptive fields were unchanged between environments providing evidence that the resolution of the egocentric location signal was robust in both familiar and novel arenas (Figure 4d; difference between baseline and novel ratemap coherence = 0.02, IQR = 0.00 – 0.08; Wilcoxon sign rank test, signed rank = 27, p = 0.25).

### Stability of EBC sub-populations requires physical boundaries

Sensory information originating from multiple modalities likely underlies the egocentric nature of the RSC boundary vector responses. There are two reasons to believe that somatosensation may inform the preferred orientation and distance of a subset of EBCs. First, many RSC neurons with egocentric boundary vector tuning had preferred distances that were proximal to the animal and within or near whisker range (Preferred distance <12cm; 16.5%, n = 21/127; Figure 1l). Secondly, the preferred orientation of EBCs spanned all egocentric bearing angles but were biased laterally perhaps reflecting whisker interaction with borders (Figure 1k). As such, we questioned whether the presence of a physical boundary was required for EBC spatial tuning and/or particular subsets of EBC receptive fields.

To this end, baseline sessions were compared to recordings in environments that were bordered by drop offs with no arena walls (n = 35 neurons from 7 sessions across 3 rats). 29 neurons recorded under these conditions exhibited EBC sensitivity in the baseline session (Figure 5g). EBCs detected in the baseline session had similar preferred orientations but more distal preferred distances in sessions with no physical walls (Figure 4b-c, difference between baseline and no walls preferred orientation = **-**6°, IQR = -70.5 – 19.5°; Wilcoxon sign rank test, z = -0.41, p = 0.68; difference between baseline and no walls preferred distance = -11.3cm, IQR = -28.8 – 2.5cm; Wilcoxon sign rank test, z = -2.4, p = 0.02). Additionally, the overall coherence of the egocentric receptive field was significantly decreased in the absence of physical walls and fewer EBCs were detected in these sessions (Figure 4d; difference between baseline and no walls ratemap coherence = 0.10, IQR = -0.00 – 0.23; Wilcoxon sign rank test, z = 3.6, p = 0.0003; EBCs with no walls = 51.4%, n = 18/35 versus 29/35). Collectively, these results suggest that the EBC population signal is degraded in the absence of explicit borders.

Despite this fact, numerous EBCs sustained their preferred egocentric vector across conditions. To investigate why some neurons were disrupted and not others we next examined the difference in receptive field coherence as a function of baseline preferred orientation and distance. There was no relationship between the preferred orientation of the neuron and the magnitude of degradation of the spatial signal with no physical walls (Figure 4h, Circular-linear correlation, ρ = 0.32, p = 0.23). In contrast, the more proximal the egocentric boundary receptive field was to the animal at baseline the more decreased the tuning was in an arena with no physical walls (Figure 4i, Spearman’s correlation, ρ = -0.39, p = 0.04). Taken together, these results support the idea that the subset of animal-proximal egocentric boundary cells (Figure 1e) may rely on somatosensory interaction with borders, while EBCs with more animal-distal receptive fields (Figure 1f) are preserved in environments with no physical walls because they rely on other sensory modalities.

### RSC EBCs are insensitive to environmental geometry which yields a directional representation of environment shape

Boundaries are unique environmental features in that they both restrict navigational affordances and define the spatial structure of the broader environment. Accordingly, the presence of boundary sensitive neurons within RSC indicates that the region is capable of detecting features of environmental geometry. In a square open field like the one utilized for baseline experimental sessions there are two primary defining features of environmental geometry: 1) conjunctions of walls forming 90° corners and, 2) boundaries that are orientated along two-axes of allocentric environmental directions. As such, we questioned if EBCs would maintain their preferred tuning in circular environments that excluded both of these geometric features.

We recorded 23 RSC EBCs as animals free foraged in square and circular environments across two experimental sessions each day (Figure 5a, total RSC neurons recorded under these conditions = 32 across 4 rats and 8 sessions). As with most other environmental manipulations, EBC boundary vectors were unchanged when the geometry of the environment was altered (Figure 5b, difference between square and circle preferred orientation = 5.5°, IQR = -33 – 18°; Wilcoxon sign rank test, z = 0.10, p = 0.92; difference between square and circle preferred distance = 2.5cm, IQR = -2.5 – 7.5cm; Wilcoxon sign rank test, z = 1.92, p = 0.054; difference between square and circle ratemap coherence = 0.002, IQR = -0.03 – 0.08; Wilcoxon sign rank test, z = 0.41, p = 0.68).

A striking feature of many EBCs (Figure 5a, **left**), but not all (Figure 5a, **right**), was the structure of movement direction tuning between square and circular environments. As a consequence of consistent egocentric boundary vector tuning in environments of different shapes, EBCs would typically possess four-pronged directional tuning that aligned with the orientation of the walls in square environments (Figure 5a, **left top**). In contrast, the same consistency in EBC tuning yielded directionally uniform tuning in circular environments (Figure 5a, **left bottom**).

Figure 5c depicts movement direction tuning plots for the full population of RSC neurons recorded in square and circular arenas. When the mean population movement direction tuning was examined, distinct peaks fell at the four cardinal directions in square arenas but no such peaks were observed in their circular counterparts (Figure 5c, **bottom plots**). We hypothesized that differences between directional tuning, as a consequence of the presence of EBCs, would allow downstream regions to disambiguate environments of different geometries.

To test this, we trained a linear classifier on a random 80% of the directional tuning curves from both environments and attempted to predict which environment the other 20% of movement direction tuning curves were recorded within (Figure 5d, Linear discriminant classifier, n = 10,000 iterations). Consistent with the hypothesis that geometry could be decoded from a population with EBC tuning, the arena could be identified correctly with 67.7% accuracy (IQR = 55.6 – 72.2%) which was statistically significant from both statistical chance (Wilcoxon sign rank with 50% accuracy median, z = 78.7, p = 0) and a classifier ran with arena identity randomized (randomized arena identity = 50%; IQR = 44.4 – 55.6%; Wilcoxon rank sum test, z = 76.6, p = 0). We conclude that regions possessing egocentric boundary vector tuning may provide punctate directional signals to downstream regions such as the medial entorhinal cortex that can be compared to other directional inputs to inform circuits about environment geometry.

### A sub-population of RSC EBCs are theta modulated

In building off of geometry detection in RSC EBC ensembles, a natural next question is how might these egocentric positional signals be integrated within the broader spatial circuity. Previous work has demonstrated that RSC local field potentials feature a prominent theta oscillation during active movement that is strongly coherent with theta rhythms observed in the dorsal hippocampal formation (Borst et al., 1987; Colom et al., 1988; Talk et al., 2004; Koike et al., 2017; Alexander et al., 2018). Spatial representations in regions with strong theta rhythmicity, such as MEC or HPC, are strongly influenced by boundaries and environmental geometry (Muller and Kubie, 1987; Gothard et al., 1996; O’Keefe and Burgess, 1996; Kenaith et al., 2017; Kinsky et al., 2018; Solstadt et al., 2008; Krupic et al., 2015; Krupic et al., 2018). We next questioned whether RSC neurons exhibiting egocentric boundary vector sensitivity were potentially synchronized with these areas via theta oscillations.

Consistent with previous work, we observed a strong RSC theta oscillation and that individual RSC neurons engage with the theta oscillation in two primary modes (Figure 6a-b). First, a small sub-population of RSC neurons exhibit theta rhythmic spiking (as revealed by autocorrelations of their spike trains) and are phase locked to the locally recorded theta oscillation (Figure 6b; 4%, n = 22/555). Second, a larger subset of RSC neurons do not possess consistent theta rhythmic spiking but are phase locked to the theta oscillation (Figure 6c; 27.6%, n = 153/555, see methods). In order to be phase modulated without firing at a theta rhythm, this latter population may transiently engage with the theta oscillation by firing at particular phases after skipping random theta cycles.

**Figure 6.**
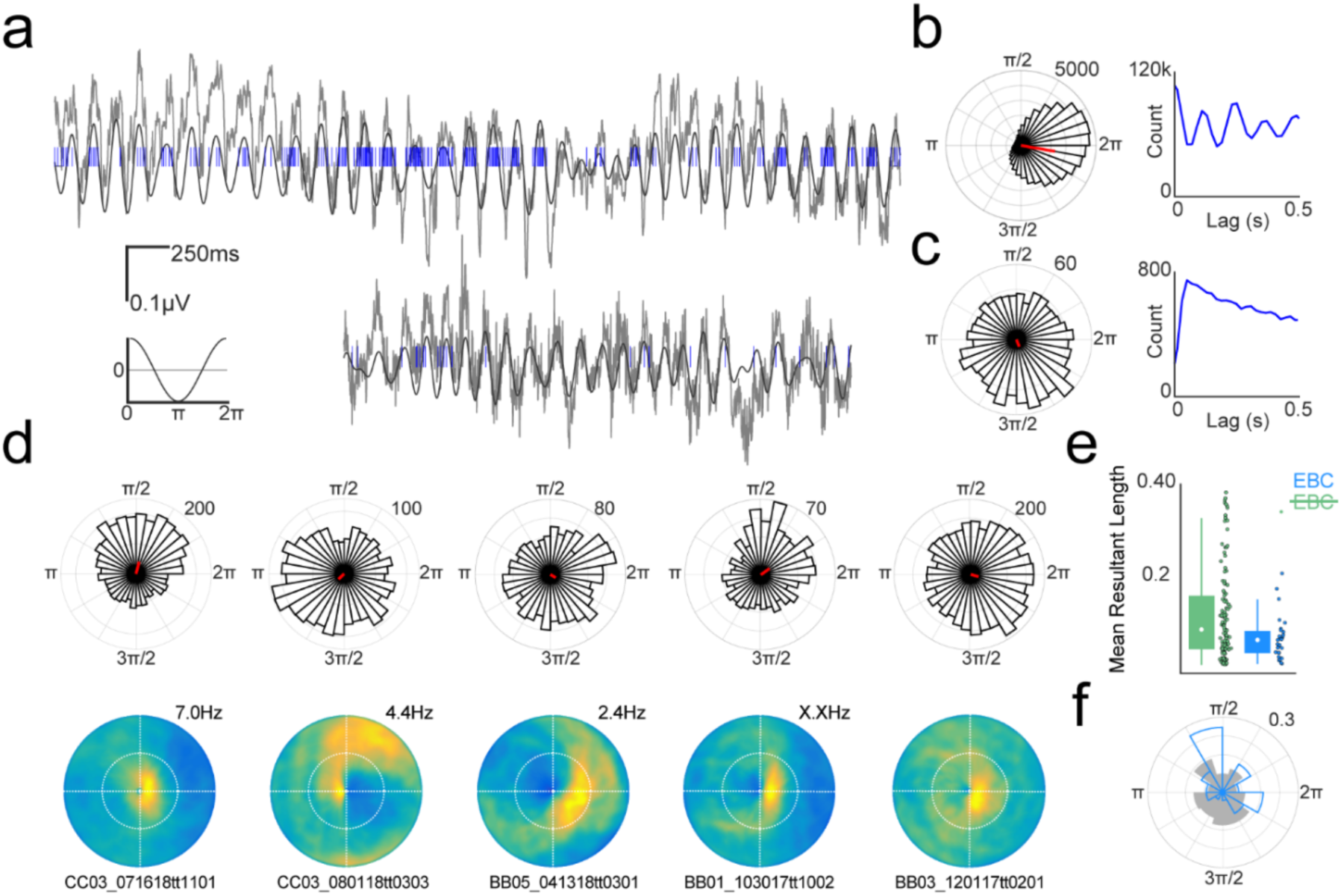
A subset of RSC EBCs are theta modulated. **a.** Two examples of RSC theta oscillation (gray) and spike train of simultaneously recorded neurons (blue). Bottom left, scale bar and schematic depicting correspondence between oscillation and theta phase. **b.** Left, circular histogram depicting spike counts as a function of theta phase for the neuron in the top row of **a**. Density of spikes near 2π indicates that the neuron is locked to the peak of the theta phase. Right, spike train autocorrelogram for the same neuron shows theta rhythmic spiking. **c**. Same as in **b,** but for the neuron depicted in the bottom row of **a**. This neuron is significantly theta phase modulated but does not exhibit theta rhythmic spiking. **d**. Example theta phase modulated EBCs. Top row, circular histogram of spike counts versus theta phase. Bottom row, corresponding egocentric boundary ratemaps. **e.** Strength of theta phase modulation as measured by the mean resultant length for non-EBCs (green) and EBCs (blue). EBCs have significantly weaker theta modulation than non-EBCs with significant phase relationships. **f.** Preferred theta phase for all EBCs (blue) and non-EBCs (gray). EBCs tended to prefer the falling phase of the theta oscillation while non-EBCs preferred the rising phase but this difference was not significant.

No EBCs exhibited intrinsically theta rhythmic spiking, but 24.4% of EBCs (n = 31/127) did phase lock to RSC theta oscillations (Figure 1d). The strength of theta modulation (mean resultant length, MRL) was significantly greater for theta-modulated non-EBCs than EBCs (Figure 6e, non-EBCs MRL = 0.13, IQR 0.10 – 0.19; EBCs MRL = 0.11, IQR = 0.09 – 0.13, Wilcoxon rank sum test, z = 2.09, p = 0.04). Although not significantly different, non-EBC theta-locked RSC neurons were biased to firing during the rising phase of the theta rhythm whereas theta-locked EBCs preferred the falling phase (Figure 6f, non-EBCs phase = 3.6 rad, IQR = 1.9 – 4.8 rad; EBCs phase = 2.3 rad, IQR = 1.7 – 5.3 rad; Kuiper two-sample test, k = 1134, p = 1). These results confirm that a sub-population of RSC EBCs are phase-locked to theta oscillations present in RSC, consistent with recent modelling work suggesting periodic modulation as a mechanism for comparing current sensory input about the environment against stored spatial representations (Byrne et al., 2007; Hasselmo et al., 2012; Bicanski and Burgess, 2018).

### RSC spatial sensitivity in the open field and during track running

RSC spatial responses have been primary examined during track running paradigms (Smith and Mizumori, 2012; Alexander and Nitz, 2015; Alexander and Nitz, 2017; Vedder et al., 2016; Mao et al., 2017; Mao et al., 2018; Miller et al., 2019). Accordingly, we questioned whether there was any relationship between stable firing correlates in free exploration and route running. To this end, a subset of RSC neurons (n=87 neurons across 3 rats) were recorded in both open field exploration and on track running paradigms (sFigure 6a).

Consistent with previous work, we observed RSC neurons with activation anchored to track locations associated with turning, start and end locations associated with reward, and non-specific patterns associated with a particular route during track running (sFigure 6b-c). Although there was no clear relationship between EBC responsivity and activation during track traversals, there was a strong positive correlation between spatial stability in these two navigational conditions, suggesting a possible sub-circuit within RSC for reliable spatial representation (sFigure 6d, open field stability (ρ) = 0.41, IQR = 0.27 - 0.60; track-running stability (ρ) = 0.07, IQR = -0.01 - 0.20; Spearman’s rho, r = 0.68, p = 0).

## Discussion

### RSC spatial representations facilitate reference frame transformations

The current data support and extend the functional role of RSC in reference frame transformations. Specifically, the RSC population exhibits sensitivity to multiple spatial coordinate systems, an essential characteristic of circuitry capable of generating such translations. In the current work we report a large subset of spatially reliable neurons that encode the position of boundaries in egocentric coordinates. Referred to as egocentric boundary cells or EBCs, these neurons robustly encoded a vectorial representation of the distance and orientation of any boundary relative to the animal itself (i.e. in an egocentric reference frame; Wang, Chen, et al., 2018; Hinman et al., 2019; LeChance et al., 2019). Egocentric boundary representations are predicted to form a critical component of the coordinate transformation circuit, as the response property could function to inform the broader spatial circuitry about the position of external landmarks in a viewpoint-dependent manner (Byrne et al., 2007; Bicanski and Burgess, 2018).

RSC neurons also exhibited multiple forms of allocentric modulation that could be integrated with EBCs or other forms of egocentric information within theta timescales. Nearly half of RSC neurons exhibited reliable and spatially-anchored responses during free foraging behavior. Spatially stable cells had complex 2D spatial representations that in some cases were reminiscent or possibly descended from spatial non-grid cells observed in mEC (Diehl et al., 2017), allocentric boundary vector cells and axis-tuned neurons of dorsal subiculum (Hartley et al. 2000; Lever et al., 2009; Olson et al., 2017), and/or location modulated head direction cells of post-subiculum (Peyrache et al., 2017). A second form of allocentric response was observed in a subset of RSC neurons that exhibit allocentric head direction sensitivity. These forms of allocentric spatial information may be processed or compared with egocentric boundary vector information within theta timescales. Both subsets of neurons exhibited theta phase modulation which is well known to synchronize information processing throughout the broader allocentric spatial circuit.

When paired with the unique anatomical connectivity of RSC with both egocentric and allocentric processing regions, the presence of neurons, such as EBCs, that are sensitive to one or more spatial coordinate systems signifies that the region is capable of interrelating external and internal spatial information for the initial construction and use of stored spatial representations. This fact may explain the diversity of impairments observed in spatial navigation, learning, and memory that occur following damage or lesion to the area (Valenstein et al., 1987; Takahasi et al., 1997; Harker and Whishaw, 2002; Vann et al., 2003; Vann and Aggleton, 2004; Vann and Aggleton, 2005; Pothuizen et al., 2008; Keene and Bucci, 2009; Hindley et al., 2014; Elduayen and Save, 2014).

### The RSC egocentric boundary vector code is context-independent which generates a directional code that reflects environment geometry

EBC spatial receptive fields were activated when the animal was positioned with both a specific orientation and distance from an environmental boundary. EBCs maintained their preferred vector tuning preference in rotated arenas, expanded arenas, and novel arenas (Figure 4). Accordingly, the EBC signal does not remap across environments, thus providing a stable, context invariant, positional metric.

This stability can be contrasted to the vast majority of allocentric representations, such as place cells, grid cells, or head direction cells, that are known to either show global or rate remapping, translations, or rotations between environments (Muller and Kubie, 1987; Bostock et al., 1991; Leutgeb et al., 2005; Yoganarasimha and Knierim, 2006; Fyhn et al., 2007; Leutgeb et al., 2007; Hoydal et al., 2019). In contrast, border cells of mEC and boundary vector cells of dorsal subiculum maintain similar tuning preferences in a context invariant manner analogous to that observed in the EBCs shown here (Solstadt et al., 2008; Lever et al., 2009). It remains to be seen what interactions exist between cells possessing these different types of boundary anchored receptive fields, however the current data suggest that boundary sensitive neurons may provide a foundational map upon which other spatial representations can be situated (Bicanski and Burgess, 2016).

Like border and boundary vector cells, RSC EBC vector representations did not remap in environments of different geometries (Solstadt et al., 2008; Lever et al., 2009). However, because EBCs respond in a directionally-dependent manner along every environmental border, the mean directional tuning of the RSC population reflected the shape of the environment (Figure 5). Here, we demonstrated that this directional signal could be utilized to distinguish arena shape. Relative positions of boundaries have repeatedly been show to alter or anchor allocentric spatial representations, especially in mEC grid cells and HPC place cells (Muller and Kubie, 1987; Gothard et al., 1996; O’Keefe and Burgess, 1996; Keinath et al., 2017; Keinath et al., 2018; Kinsky et al., 2018; Solstadt et al., 2008; Krupic et al., 2015; Krupic et al., 2018; Julian et al., 2018). mEC receives excitatory projections, both directly and indirectly, from RSC and projects into HPC (Kononenko and Witter, 2012; Czajkowski et al., 2013). We hypothesize that the RSC arena-geometry-related directional signal may serve to provide excitatory drive at specific allocentric head directions to inform the circuit about the relative angles amongst borders.

### Egocentric vector tuning may support route-centric representations of RSC and PPC

Consistent with previous work in dorsal striatum, RSC EBC tuning was strongest during motion and the directional component of egocentric boundary receptive fields was more robustly driven by the movement direction of the animal rather than head direction (Hinman et al., 2019). Taken together, these properties of EBC sensitivity suggest that the positional signal is related to active navigation and the relationship of trajectories through the environment relative to environmental boundaries.

RSC and the reciprocally connected PPC have been shown to exhibit activity patterns during track running paradigms that are anchored to the shape of the route itself (Nitz, 2006; Nitz, 2012; Alexander and Nitz, 2015; Alexander and Nitz, 2017). Route-referenced activity in these regions could be potentially explained by EBC tuning or in part arise from the integration of EBC responsivity with other spatial covariates. Consistent with this hypothesis, we showed that the sub-population of spatially stable neurons in 2D free foraging (which included large numbers of EBCs) were more likely to exhibit spatially stable representations during track running (sFigure 6).

### EBCs are primarily restricted to the dysgranular RSC

A striking anatomical feature of the EBC population was that it was primarily localized to the dysgranular sub-region of RSC (dRSC). dRSC has connectivity weighted towards egocentric coordinate systems, as it is reciprocally innervated by cortical regions important for processing sensory and motor information as well as association areas such as PPC wherein egocentrically-referenced spatial responses have been observed (McNaughton et al., 1994; Whitlock et al., 2012; Wilber et al., 2014; Wilber et al., 2017). Further, the concentration of EBCs in dRSC is consistent with theoretical work posing a circuit for translating between egocentric and allocentric coordinate systems that includes posterior parietal cortex (PPC), RSC, and the extended hippocampal formation as primary hubs (Byrne et al., 2007; Oess et al., 2017; Rounds et al., 2018; Clark et al., 2018; Bicanski and Burgess, 2018).

Of note, dRSC is also known to possess bidirectional head direction cells (BDHD) that respond to local reference frames in multi-compartment environments with distinct contextual cues (Jacob et al., 2017). This sensitivity ultimately yields allocentric directional tuning plots that are bimodal. In the current work, strongly tuned EBCs commonly exhibited quad-modal allocentric directional tuning that was aligned with the four walls of square environments. This similarity in directional tuning response of EBCs and BDHDs and their co-localization in dRSC raises questions as to the nature of the relationship or interactions between these functional sub-populations.

One possibility is that neurons in dRSC are prone to represent the locations of spatial landmarks using egocentric vectors and that EBCs and BDHDs are both special cases constrained by their respective experimental setups. In the case of EBCs reported here, the vector may anchor to boundaries because borders are the only landmarks present in the open field that can cause activation of the receptive field. In the work of Jacob et al., the egocentric vector may respond to borders as well as local visual landmarks or doorways between two compartments. The bimodal directional tuning in the latter experiment may arise from constrained egocentric sampling along two-axes as a consequence of the multi-compartment environment segmenting two opposing walls. This proposed egocentric vector encoding of environment features in RSC may underlie functional correlates of local heading orientation, scene processing, or goal location in RSC in humans (Maguire, 2001; Epstein et al., 2007; Epstein et al., 2008; Marchette et al., 2014; Chrastil et al., 2015; Patai et al., 2019).

### A network of vector-based egocentric spatial representation

In addition to RSC, EBCs were also observed in both posterior parietal (PPC) and secondary motor cortices (M2) but not mEC which is commonly thought to represent space in allocentric coordinates. The presence of EBCs in PPC converges nicely with previous work demonstrating egocentric bearing sensitivity of PPC neurons to visual cues positioned along boundaries (Wilber et al., 2014). Computational models exploring circuitry for reference frame transformations and spatial imagery initially predicted EBCs to exist in PPC (Byrne et al., 2007). However, egocentric responses were initially reported in lateral entorhinal cortex, dorsal striatum, and postrhinal cortex, and now here in RSC, PPC, and M2 (Wang, Chen, et al., 2018; Hinman et al., 2019; LeChance et al., 2019; see also, Gofman et al., 2017). Accordingly, a picture of a distributed network of interconnected regions with egocentric vector representations is beginning to emerge. Given the presence of EBCs in several midline structures, it is possible that EBCs are also present in the anterior cingulate cortex as well as thalamic structures that innervate midline associative cortex (Weible et al., 2012; Matulewicz et al., 2019). Future investigations should focus on dependencies amongst the regions currently implicated, as the EBC network may possess functional and anatomical connectivity resembling the well-characterized extended head direction cell network (Taube et al., 2007).

## Acknowledgements

We would like to thank Madelynne Campbell, Paola Castro-Mendoza, Jordan Dreher, Elin Johansson, Jongyoul Lee, and Pedro Rodriguez-Echemendia for technical assistance. We also thank Jacob Olson and Douglas Nitz for useful discussions and comments pertaining to the manuscript. This research was supported by National Institutes of Health NINDS F32 NS101836-01, NIMH R01 MH061492 and MH060013, and Office of Naval Research MURI grant N00014-16-1-2832.

## Author Contributions

A.S.A., L.C.C., J.R.H., and M.E.H designed the study. A.S.A and L.C.C. conducted all experiments. A.S.A., F.R., and G.W.C. analyzed the data. A.S.A. and M.E.H wrote the paper. All authors assisted with revision of the manuscript.

## Declaration of Interests

The authors declare no competing interests.

## Supplemental Figures

**Supplemental Figure 1.**
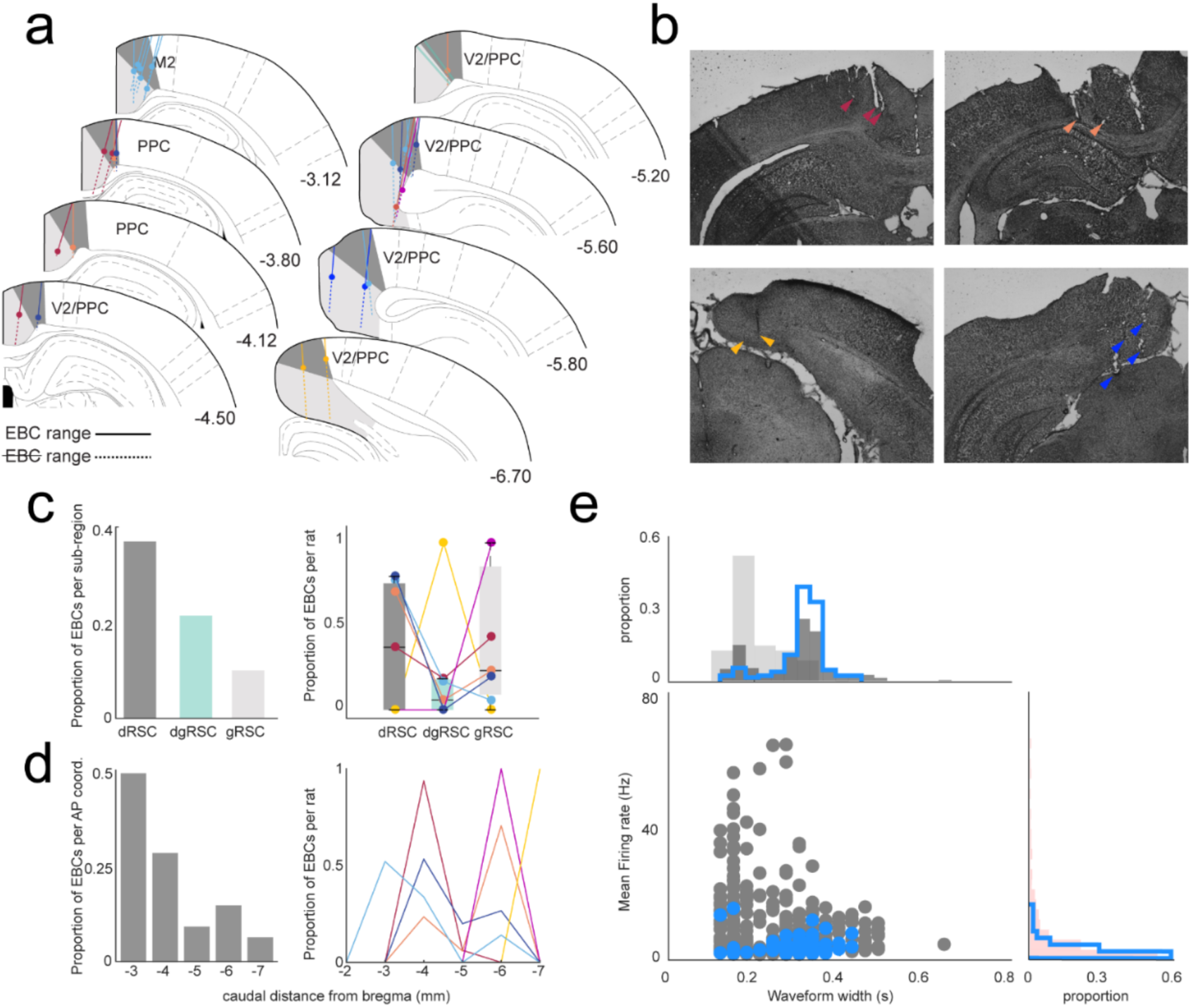
Locations of RSC egocentric boundary cells and identification as putative principal cells. **a.**Schematic depicting location of tetrodes where EBCs were recorded. Each color corresponds to a different animal. Dark gray indicates dRSC and light gray indicates gRSC. Circles indicate the approximate dorsal-ventral (D/V) position of the wire for the last recording with EBCs on each tetrode. The dashed line indicates span of recordings in D/V axis that did not yield EBCs. Solid line indicates span of recordings in D/V axis that did yield EBCs. EBCs were also observed in secondary motor cortex (M2) or parietal cortex (V2/PPC) and these regions are noted. **b.** Example histology showing tetrodes in RSC. Tetrode locations are indicated with colored triangles which correspond to individual animals in **a**. **c.** Left, proportion (out of 1.0) of EBCs relative to all RSC neurons recorded in each RSC sub-region. Right, proportion of EBCs relative to all RSC neurons recorded in each sub-region for each animal (colored dots and lines). **d.** Same as for **c**, but for approximate anterior-posterior location of EBC. **e.** Bottom left, scatterplot depicting mean firing rate versus width of spike waveform for all RSC neurons in gray and EBCs in blue. Top, histogram of waveform width clustered into fast (light gray) and slow waveforms (dark gray). Blue histogram corresponds to distribution of waveform width for all EBCs which have wider waveform widths. Right, histogram of mean firing rate (light pink). Blue histogram corresponds to distribution of mean firing rate of all EBCs demonstrating overall low mean firing rate.

**Supplemental Figure 2.**
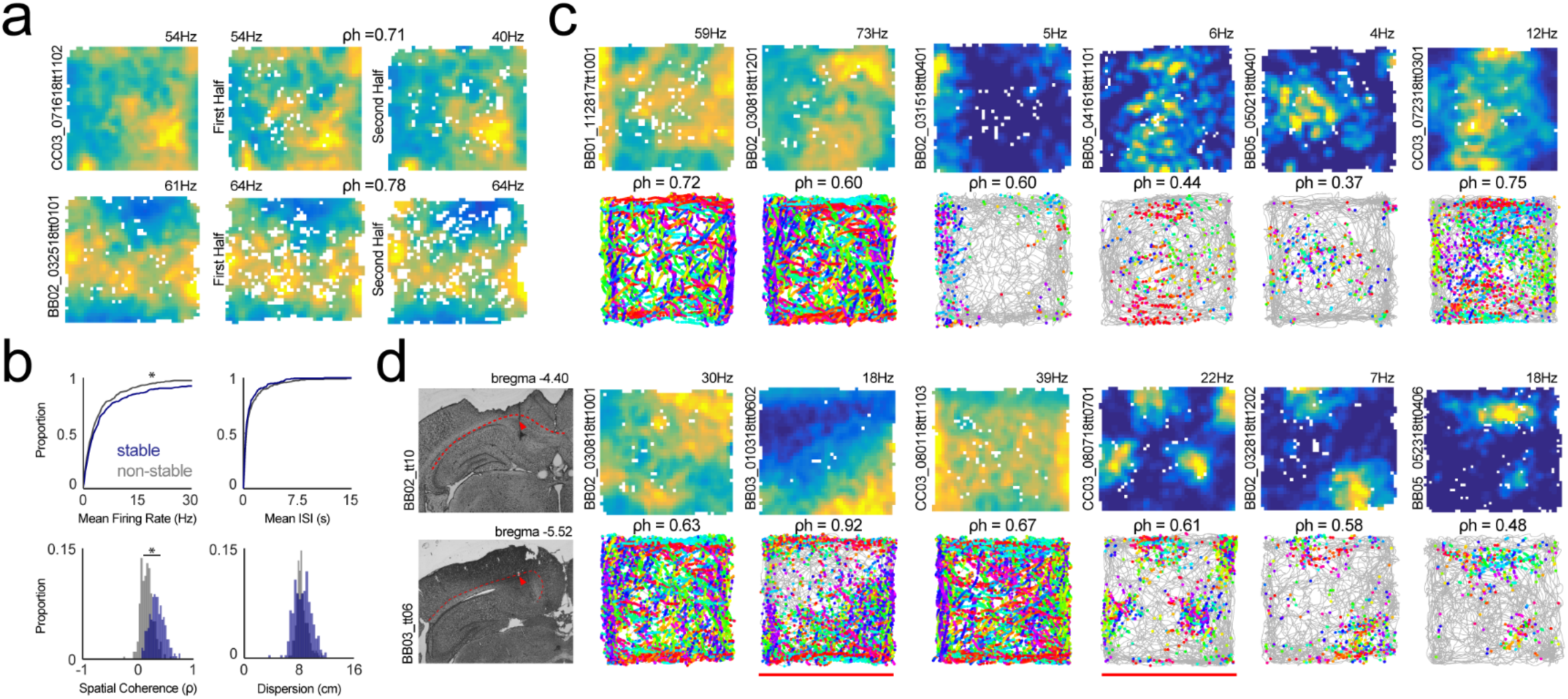
RSC spatial stability during free foraging. **a.**Two-dimensional (2D) firing ratemaps for two neurons (rows) showing process for determining spatial stability. Left column, 2D ratemap for full session. Middle and right columns, 2D ratemaps for the first and second halves of each session. Above ratemaps, corresponding spatial correlations between the two halves of the session. **b.** Properties of spatially stable (blue) versus non-stable (gray) RSC neurons. Top left, spatially stable cells had slightly lower mean firing rates than those that did not have spatial reliability (sFigure2b; stable = 3.02, IQR = 1.23 - 7.25Hz; not-stable = 4.93, IQR =0.95 - 5.56Hz; Wilcoxon rank sum, z= -1.06, p = 0.04). Top right, mean inter spike intervals were not different between stable and non-stable neurons (stable = 0.36, interquartile range (IQR) = 0.15 - 0.99; non-stable = 0.41, IQR = 0.19 - 1.15; Wilcoxon rank sum, z = 1.89, p = 0.06). Bottom left, RSC neurons with spatially reliable activity had significantly greater spatial coherence than non-stable cells (stable Spearman’s ρ = 0.33, IQR = 0.25 - 0.43; not-stable Spearman’s ρ = 0.16, IQR =0.08 - 0.23; Wilcoxon rank sum, z= -14.448, p = 2.6×10^−47^). Bottom right, spatial dispersion of the top 90% of firing rate bins in centimeters for stable and non-stable neurons (stable = 21.3cm, interquartile range (IQR) = 18.8 - 23.3cm; non-stable = 20.6cm, IQR = 19.7 - 22.2cm; Wilcoxon rank sum, z = 1.89, p = 0.06). **c**. 2D ratemaps and trajectory plots for example RSC neurons with significant spatial stability. **d.** Example 2D firing rate maps and trajectory plots for neurons recorded at or near the cingulum bundle. Left column shows two such recording locations with neurons recorded at each site demarcated with a horizontal red line. In some cases, recordings from the cingulum bundle resembled grid cells.

**Supplemental Figure 3.**
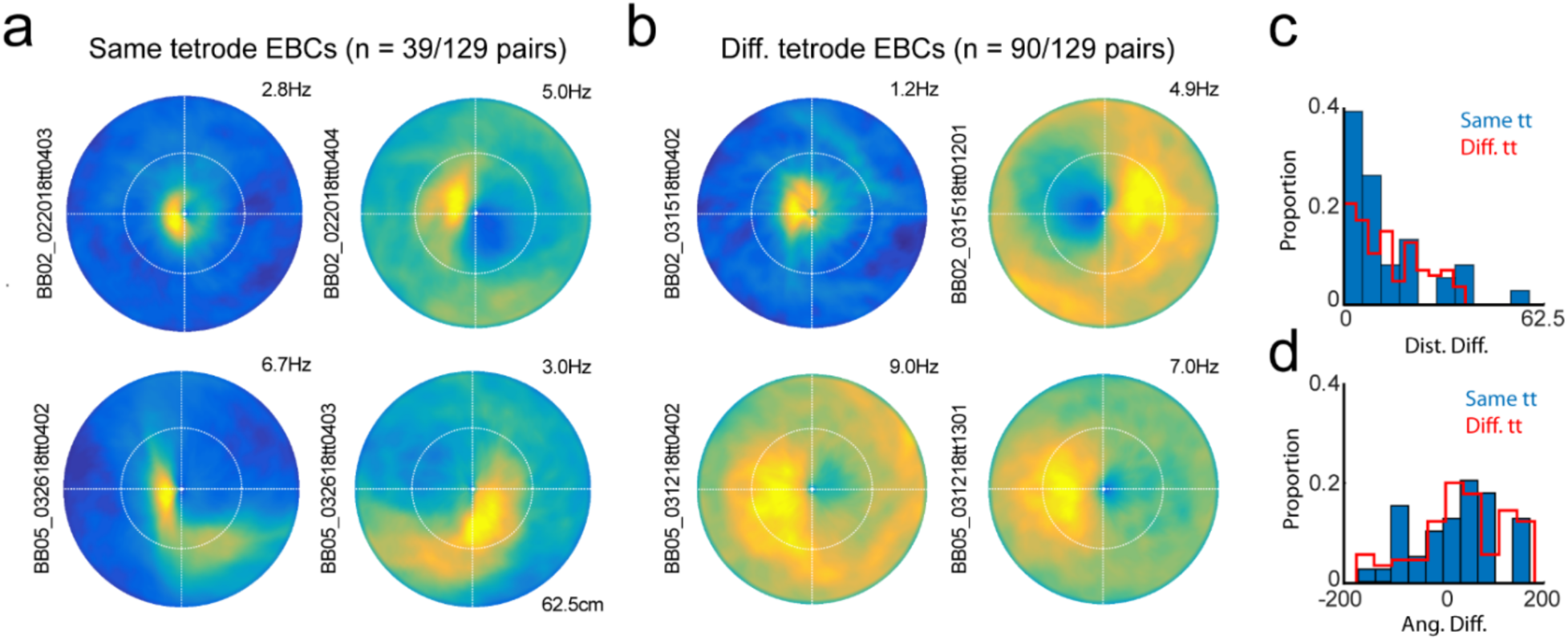
Simultaneously recorded egocentric boundary cells. **a.**Each row depicts egocentric boundary ratemaps for a pair of simultaneously recorded egocentric boundary cells that were observed on the same recording tetrode. Top pair have similar EBC receptive field locations while bottom pair have receptive fields on different sides of the animal. **b.** Same as in **a**, but for simultaneously recorded neurons on different tetrodes. Top pair have different receptive field locations while bottom pair have similar receptive fields. **c.** Histogram of absolute difference in preferred distance for pairs of simultaneously recorded EBCs on the same tetrode (in blue) and on different tetrodes (in red). **d.** Histogram of difference in preferred orientation for pairs of simultaneously recorded EBCs on the same tetrode (in blue) and on different tetrodes (in red).

**Supplemental Figure 4.**
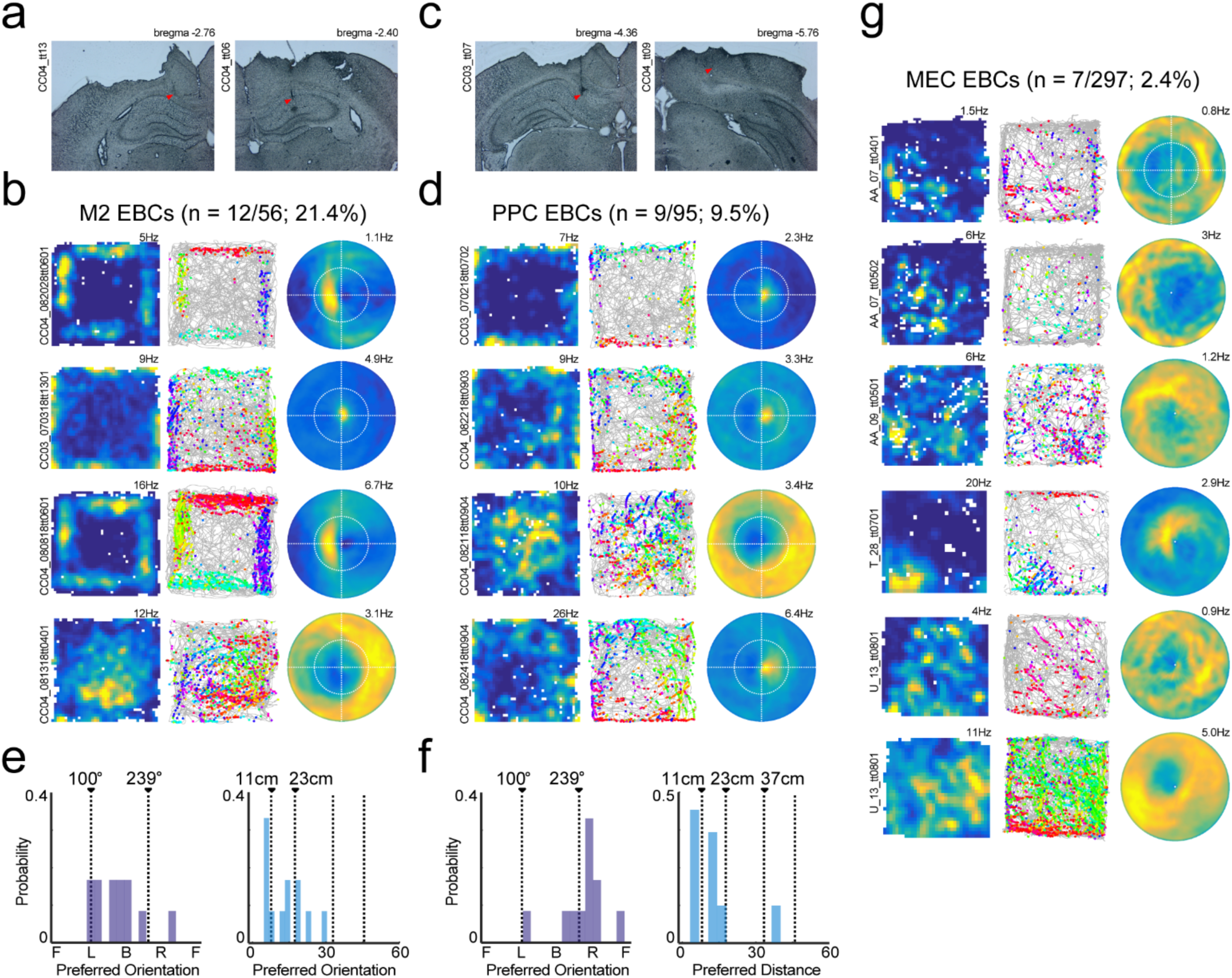
Egocentric boundary cells in secondary motor cortex and posterior parietal cortex but not medial entorhinal cortex. a. **a.** Example histology showing placement of recording tetrodes in secondary motor cortex (M2). **b.** 2D ratemaps, trajectory plots and egocentric boundary ratemaps (EBR) for 4 example M2 neurons with EBC sensitivity. **b** Example histology showing placement of recording tetrodes in parietal cortex (V2/PPC). **d.** 2D ratemaps, trajectory plots and egocentric boundary ratemaps (EBR) for 4 example V2/PPC neurons with EBC sensitivity. **e.** Preferred orientation and distance for M2 neurons identified as EBCs. **f.** Preferred orientation and distance for V2/PPC neurons identified as EBCs. **g.** All detected medial entorhinal cortex EBCs.

**Supplemental Figure 5.**
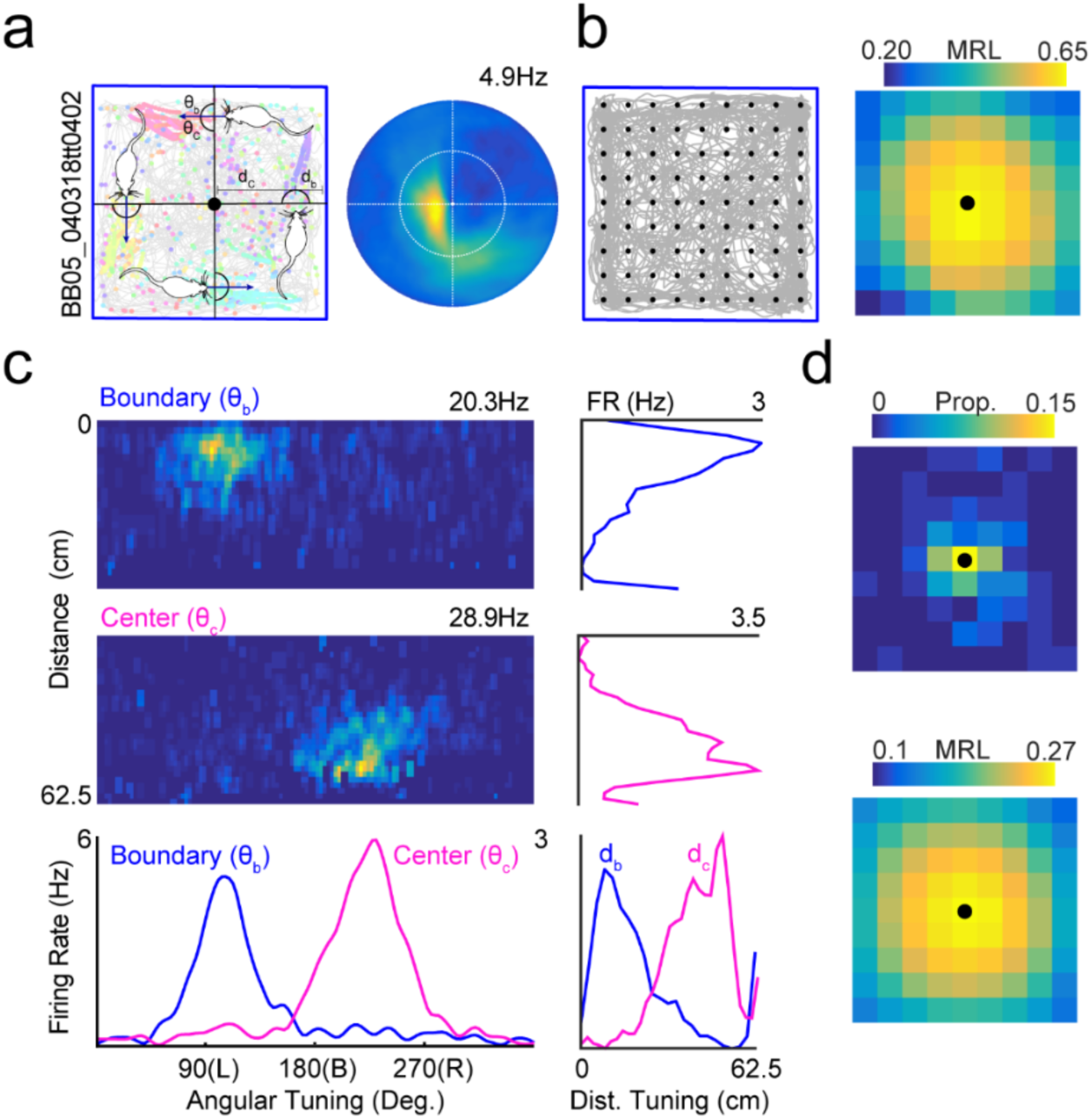
Egocentric vector tuning to center of arena for generalized linear models. **a.** Left, schematic depiction of the angular and distance relationships between the animal, the boundary, and the arena center when the animal is running parallel to arena walls. Right, egocentric boundary ratemap for an EBC with trajectory plot on left plot. **b.** To analyze whether the egocentric bearing and distance to the arena center could function as a predictor in generalized linear modelling we analyzed strength of egocentric bearing and distance tuning to a 9×9 grid of locations that spanned the entirety of the environment (see Wang, Chen, et al., 2018). Right plot depicts the strength (as measured by mean resultant length, MRL) of egocentric vector tuning to all 9×9 grid locations for the neuron depicted in **a.** The heatmap corresponds to magnitude of the MRL, and the black dot depicts the grid location with the strongest egocentric tuning which, in this case, corresponds to the center of the arena. **c.** For the same neuron we calculated egocentric bearing and distance tuning to the nearest point on the boundary (top left EBC heatmap) and the grid location with maximum tuning (center of arena, bottom left EBC heatmap), and compared their vector components by averaging across angular and distance dimensions of each matrices. Plots to the right of each EBC heatmap show average distance tuning to the nearest boundary (blue) and center (pink) with each overlaid on the bottom right plot. Below the EBC heatmaps are overlaid averages of preferred angular tuning for nearest boundary (blue) and arena center (pink). The tuning plots at the bottom for both reference points show a clear relationship and so we utilized the egocentric boundary and distance to the central location for GLM analyses. **d.** Top, 2D histogram showing the proportion of RSC EBCs with maximum MRLs at each grid location. Greatest proportion is at or near the center of the arena in baseline sessions when boundaries are excluded. Bottom, mean MRL tuning for all grid locations across all RSC EBCs shows maximum egocentric vector tuning to the arena center in baseline sessions when boundaries are excluded.

**Supplemental Figure 6.**
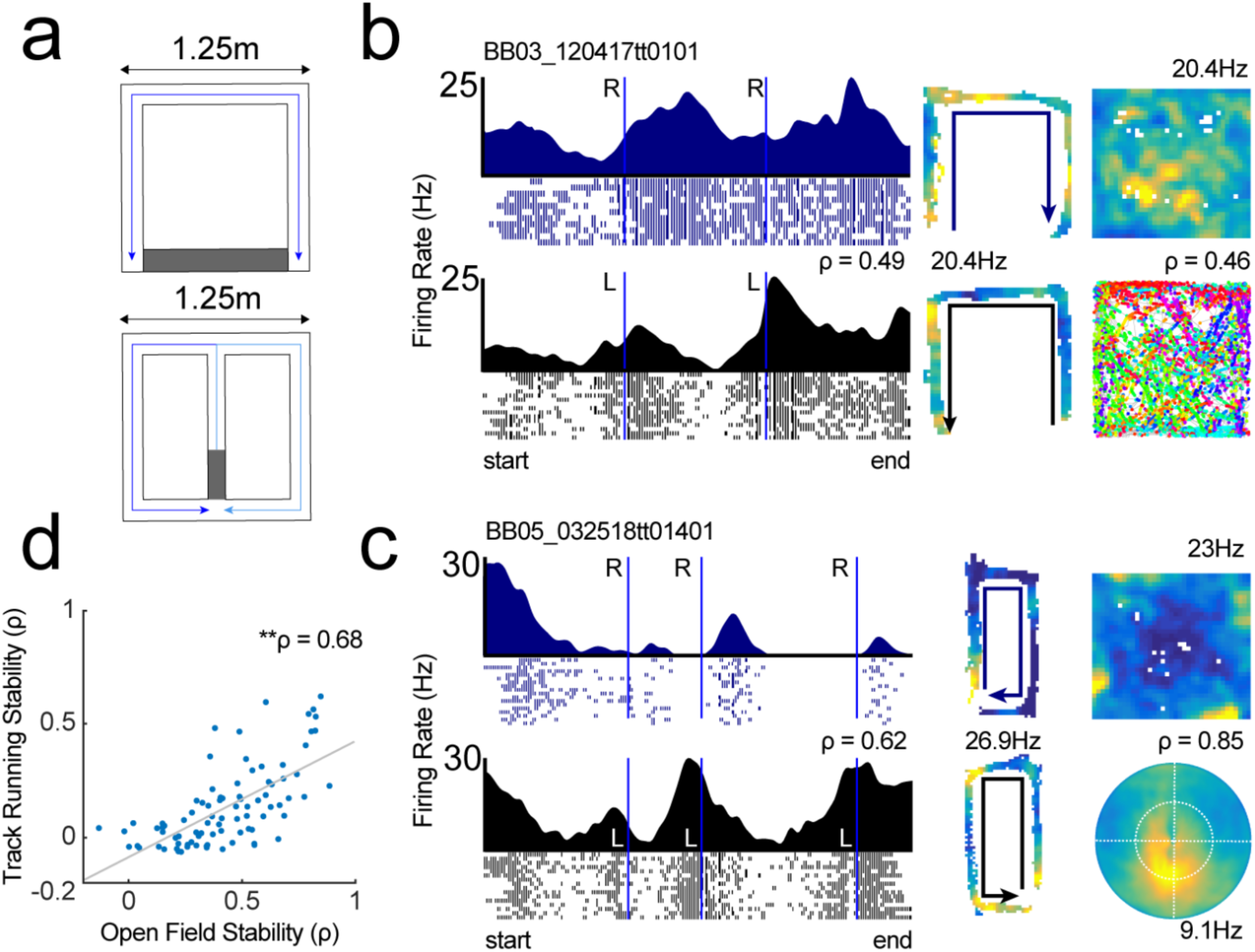
Comparison between RSC neurons recorded in free foraging and during track running. **a.** Schematic of two different tracks animals ran along. Gray zone indicates reward location. **b**. Linearized firing rate vectors split into left-turning and right-turning routes with corresponding 2D ratemaps for two RSC neurons. Left column in blue, mean firing rate for route with right turns and trial spike trains. Left column in black, mean firing rate for route with left turns and trial spike trains. 2D ratemaps for track running depicted in middle column. Right column, 2D ratemap and trajectory plot. **c.** Same as **b**, but right column depicts 2D ratemap and corresponding egocentric boundary ratemap. This neuron shows spatial selectivity on track near turns and is an egocentric boundary neuron in the open field. Critically, the EBC receptive field does not explain apparent counterclockwise (left) turning preference on the track. **d.** Scatterplot depicting Spearman’s ρ for open field stability versus track running stability for all RSC neurons recorded in both conditions. Significant positive correlation indicates that neurons that were stable in one condition were more likely to be stable in the other.

## Methods

### Subjects

Male Long-Evans rats (n = 7; Charles River Labs, Wilmington, MA) were housed individually in plexiglass cages and kept on a 12-h light/dark cycle. Rats had continuous access to food during a habituation period lasting approximately 1 week. After this period, animals were food restricted until they reached 85-95% of their weight during free feeding. Water was available continuously. All procedures were approved by the Institutional Animal Care and Use Committee at Boston University.

### Shaping/behaviour

Animals were acclimated to the primary testing room for approximately one week. During acclimation, rats were handled by multiple researchers and trained to consume both Froot Loops (General Mills, Battle Creek, MI) and 45mg chocolate pellets (Bio-Serv, Flemington, NJ). After animals readily ate both food items they were exposed to one of two familiar open fields used for baseline sessions for 20 to 45 minutes per day. The first open field was 1.25m^2^ with four black walls 30 cm in height. The second arena was 1.25m^2^ with three black walls and one white wall 30 cm in height. Both arenas were placed on a dark gray textured rubber floor that was cleaned between sessions. Two animals (BB01 and BB03) performed a goal directed navigation task in a different arena and testing room prior to being utilized for the current study.

### Surgical procedures

Rats were surgically implanted with custom-fabricated hyperdrives in aseptic conditions. Each hyperdrive was composed of 12 to 16 nickel chromium tetrodes (12µm, Kanthal-Sandvik, Hallstahammar, Sweden) that could be independently moved in as small as 35µm increments. Guide cannulae for each tetrode were collectively configured in one of three arrays: 1) filling a single hypodermic tube approximately 2mm^2^ in diameter, 2) across two conjoined hypodermic tubes that were ∼1.5mm^2^ in diameter spanning a total of ∼3mm or, 3) across four conjoined hypodermic tubes that were ∼1.25mm^2^ in diameter and spanned a total of ∼5mm. For the second and third configurations, the long axis of the electrode array was positioned to target an extended region of the anterior-posterior axis of retrosplenial cortex.

Animals were anesthetized using a combination of inhaled isoflurane (0.5% initial concentration) and ketamine/xylazine administered via intraperitoneal injection (Ketamine: 12.92 mg/kg, Acepromazine: 0.1mg/kg, Xylazine: 1.31 mg/kg). After the animal was determined to be under anesthesia (as assessed via loss of the toe pinch reflex), the animal was positioned in a stereotaxic apparatus, a 0.1mg/kg dose of atropine was administered subcutaneously, and the head was shaved. Excess hair was removed via application of Nair (Church & Dwight Co., Ewing, NJ) and the scalp was cleaned with 70% ethanol and Betadine (Avrio Health L.P., Stamford, CT). 0.9% sodium chloride was administered subcutaneously hourly throughout the surgical procedure.

Following a midline incision and subsequent clearing of connective tissue, a ground screw was positioned above the cerebellum and 5-8 anchor screws were affixed in an array around the perimeter of the exposed skull. A large craniotomy was centered above retrosplenial cortex (relative to bregma: A/P: -2mm to –7mm; M/L ±0mm-1.75mm). The exact size and position of the craniotomy was dependent upon the aforementioned configuration of the hyperdrive array. Next, dura was resected and the hyperdrive was positioned such that guide cannula rested gently against the dorsal surface of the brain. Excess exposed tissue within the craniotomy was protected with Kwik-Sil (World Precision Instruments, Sarasota, Fl), and the implant was secured to anchor screws with dental cement. Tissue around the implant was cleaned with saline, 70% ethanol, and hydrogen peroxide. Antibiotic ointment was applied into the wound, sutured if necessary, and Neosporin was applied around the site. Prior to removal from anesthesia tetrodes were lowered approximately 0.25mm D/V. Animals received post-operative antibiotics (Baytril: 10mg/kg) and analgesics (Ketofen: 5.0mg/kg) for five days after surgery and were freely fed. After one-week post-operation animals were handled and reacclimated to the testing room and free foraging environments prior to the initiation of experiments.

### Electrophysiological recordings

Neural signals were amplified at two headstages attached to a 64 channel electrical interface board and acquired by a 64 channel Digital Lynx acquisition system (Neuralynx, Bozeman, MT) Signals were digitized, filtered (0.3-6.0kHz), and amplified (5,000-20,000X). Timestamps of individual action potentials were detected online when the signal crossed an acquisition threshold on any individual electrode composing a tetrode. At the conclusion of each experiment, spikes were manually sorted to individual single units using Offline Sorter (Plexon Inc., Dallas, TX) and the following features: peak-valley, peak, and principal components 1-3. Two diodes attached to the electrode implant delineated the location of the animal which was tracked at 30Hz via a camera positioned above the recording arena.

An experimental session began with an initial 15-30 minute recording while the animal free explored and consumed either Froot Loops scattered by an experimenter, chocolate pellets released at random intervals from a dispenser positioned above the arena, or both (Open Ephys Pellet Dispenser designed by Maurer Lab, https://github.com/jackpkenn/PelletDispenser). After adequate spatial coverage was achieved the animal was removed from the arena and placed back in its home cage for a period of 1 hour minimally. During this period, the experimenter completed clustering of action potentials into single neurons and examined two-dimensional spatial ratemaps (described below) to assess whether any of the cells from the baseline session exhibited egocentric boundary vector sensitivity. If no EBCs were present, tetrodes were typically lowered between 35 and 70µm. If EBCs were present, a second experimental session was conducted in which the animal explored an open field in one or more of the following configurations:

1. Open field session: The same arena from the baseline session to assess the stability of EBCs in familiar environments.
2. Open field rotation: The same arena from the baseline session rotated 45° relative to the testing room and all visible distal cues present therein.
3. Circular open field: A familiar circular arena of 1.1m diameter.
4. Open field expansion or contraction: If an expansion experiment was planned the initial baseline session was conducted in a familiar 1.25m^2^ arena that enabled reconfiguration of walls. Following the baseline session, walls were uniformly moved outwards relative to the center point of the baseline configuration to a size of 1.5m^2^ or larger. In a small number of recordings, walls were either: moved non-uniformly to increase the length of the arena along a single axis of the environment or, contracted to decrease the size of the environment. Across all possible wall movements, the arena was altered in size by approximately 20% along each dimension.
5. No wall open field: If a no wall arena experiment was planned the initial baseline session was conducted in a familiar 1.25m^2^ environment that was placed approximately 20cm above the floor. Following this session, the walls were removed from this arena creating a platform with no walls that the rat explored for a second time. In a small number of sessions, the animal explored the familiar 1.25m^2^ that was situated on the testing room floor, then in a second session, explored a different familiar arena lacking walls placed approximately 8” above the floor.

All arenas were positioned such that the animal could easily see the broader recording room and an array of stable distal cues. In some cases, the manipulation session was followed by a return session to the familiar baseline arena.

### Histology

Animals were anesthetized with 0.5% isoflurane and small electrical lesions were made at the end of tetrodes that had preliminarily been identified as having EBCs. After one week, animals were deeply anesthetized with isoflurane, injected with sodium pentobarbital, and transcardially-perfused with 0.9% saline followed by 10% formalin. The brain was extracted from the skull and post-fixed overnight with 10% formalin, then stored in 0.1M phosphate buffer until two days before slicing when it was transferred to a 0.1M phosphate buffer/30% sucrose solution. The brain was snap frozen using 2-methylbutane and sliced into 40-50um coronal sections using a cryostat (Leica CM3050S, Leica Biosystems, Buffalo Grove, IL). Slices were mounted on gelatin covered microscope slides and allowed to dry, then photographed (Nikon DXM1200 camera mounted on Olympus BX51 light microscope). Tetrode lesions and tracts were clearly visible in all animals. Coordinates of tetrode locations and final tetrode depths were registered with respect to pre-implant photographs of guide cannulae array configurations and tetrode turning logs, respectively.

### Data Analysis

#### Two-dimensional (2D) spatial ratemaps and spatial stability.

Animal positional occupancy within an open field was discretized into 3×3cm spatial bins. For each neuron, the raw firing rate for each spatial bin was calculated by dividing the number of spikes that occurred in a given bin by the amount of time the animal occupied that bin. Raw firing rate maps were smoothed with a 2D Gaussian kernel spanning 3cm to generate final rate maps for visualization. Individual raw firing rate maps were also computed after dividing the session into halves. To assess spatial stability of an individual RSC neuron, the similarity of the two raw firing rate maps from non-overlapping halves of the recording session was calculated using the non-parametric Spearman’s rank correlation coefficient. To determine whether a given spatial stability value was greater than expected by chance we next conducted randomization tests wherein the spike train for each RSC neuron was circularly shifted relative to spatial position 100 times, and individual firing rate maps were constructed for non-overlapping halves that were then correlated. The spatial stability correlation values following randomizations were collapsed into a single distribution for all neurons and randomizations and the 99^th^ percentile of all values was calculated. RSC neurons with real spatial stability correlations greater than this threshold were determined to have robust two-dimensional spatial stability.

#### Construction of egocentric boundary ratemaps

Egocentric boundary ratemaps (EBR) were computed in a similar manner as 2D spatial ratemaps but referenced relative to the animal rather than the spatial environment. The position of the boundaries relative to the animal was calculated for each position sample (i.e. frame). For each frame, we found the distance, in 2.5cm bins, between arena boundaries and angles radiating from 0° to 360° in 3° bins relative to the rat’s position. Critically, angular bins were referenced to the current movement direction of the animal such that 0°/360° was always directly in front of the animal, 90° to its left, 180° directly behind it, and 270° to its right. Intersections between each angle and environmental boundaries were only considered if the distance to intersection was less than or equal to ½ the length of the most distant possible boundary (in most cases this threshold was set at 62.5cm or half the width of the arena). In any frame the animal occupied a specific distance and angle relative to multiple locations along the arena boundaries, and accordingly, for each frame, the presence of multiple boundary locations were added to multiple 3° x 2.5cm bins in the egocentric boundary occupancy map. The same process was completed with the locations of individual spikes from each neuron, and an EBR was constructed by dividing the number of spikes in each 3° x 2.5cm bin by the amount of time that bin was occupied in seconds. Smoothed EBRs were calculated by convolving each raw EBR with a 2D Gaussian kernel (5 bin width, 5 bin standard deviation).

For EBR construction and other analyses in the current work, movement direction was defined as the instantaneous derivative of the position signal whereas head direction was the instantaneous angle calculated from the location of two position tracking diodes. Movement direction was utilized as the primary directional variable in EBR construction but a comparison to head direction determined the latter to be a less robust signal for egocentric boundary vector tuning.

#### Head direction cell identification

For each neuron, the mean resultant length of the firing rate as a function of head direction was calculated as

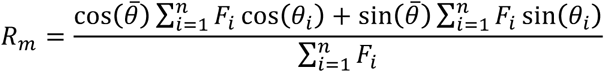

where 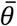 was the head direction of firing and *F_i_* and θ_*i*_ were the firing rate and head direction for bin *i*. Head direction cells (HD) were identified as those cells with *R_m_* greater than 0.20. HD cells (n = 27/555; 4.9% of all RSC neurons) were removed from the possible pool of RSC EBCs.

#### Identification of neurons with egocentric boundary vector tuning

To identify neurons with significant egocentric boundary vector sensitivity we began by calculating the mean resultant (MR) of the cell’s egocentric boundary directional firing collapsed across distance to the boundary. The mean resultant was calculated as

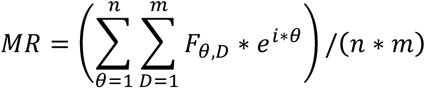

where θ is the orientation relative to the rat, *D* is the distance from the rat, *F_θ,D_* is the firing rate in a given orientation-by-distance bin, *n* is the number of orientation bins, *m* is the number of distance bins, *e* is the Euler constant and *i* is the imaginary constant. The MRL is defined as the absolute value of the mean resultant and characterized the strength of egocentric bearing tuning to environment boundaries. We next computed the preferred orientation of the egocentric boundary ratemap as the mean resultant angle (MRA)

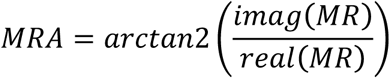

We estimated the preferred distance by fitting a Weibull distribution to the firing rate vector corresponding to the MRA and finding the distance bin with the maximum firing rate. The MRL, MRA, and preferred distance were calculated for each neuron for the two halves of the experimental session independently. Next, the MRL was computed for each neuron following 100 random, unrestricted, circular shifts of the spike train relative to position. The 99^th^ percentile of the MRL distribution across all neurons was determined.

A neuron was characterized as having egocentric boundary vector tuning (i.e. an EBC) if it reached the following criteria: 1) the MRL for both halves of the baseline session were greater than the 99^th^ percentile of the randomized distribution, 2) the absolute circular distance in preferred angle between the 1^st^ and 2^nd^ halves of the baseline session was less than 45°, and 3) the absolute difference in preferred distance between the 1^st^ and 2^nd^ halves of the baseline session was less than 75% of the preferred distance for the entire session.

To refine our estimate of the preferred orientation and preferred distance of each neuron we calculated the center of mass (COM) of the receptive field defined after thresholding the entire EBR at 75% of the peak firing and finding the largest continuous contour (‘contour’ in Matlab). We repeated the same process for the inverse EBR for all cells to identify both an excitatory and inhibitory receptive field and corresponding preferred orientation and distance for each neuron.

#### Clustering of EBC receptive field sub-types

In an effort to identify trends in receptive field sub-types including those exhibiting inverse receptive fields, we next clustered the EBC sub-population using k-means (‘kmeans’ in MATLAB) and the following feature space: distance from the origin to the global minimum of the EBR, distance from the origin to the global maximum of the EBR, the stepwise percentage of the EBR that was greater than 20% to 90% of the maximum firing rate in 10% bounds, and the dispersion of the EBR and inverse EBR. All features were z-score normalized across the population of EBCs prior to clustering. K-means was run on these features for up to 10 clusters and the total within-cluster sums of point-to-centroid distances (SUMD) were examined to assess which K was appropriate for the current data. Repeated iterations of k-means clustering and qualitative inspection of the K vs SUMD plot revealed a consistent elbow at K = 4 clusters. Principal components analysis (‘pca’, in MATLAB) was run on the same features to visualize whether k-means was partitioning distinct clusters or a continuum.

#### Ratemap coherence, dispersion, and receptive field size

For either egocentric boundary ratemaps or 2D spatial ratemaps, coherence was defined as the Spearman’s correlation between each spatial bin and the mean firing rate of all adjacent bins. Dispersion was calculated as the mean within rate map distance of the top 10% of firing rate bins. Receptive field size was only calculated for egocentric boundary ratemaps (described below), and was defined by the area (percentage of all egocentric boundary ratemap degree x cm bins) of the largest single contour detected after selecting for bins with firing rates greater than 75% of the maximum firing rate.

#### Self-motion rate maps and assessment of self-motion sensitivity

Angular displacement (θ) was calculated by determining the circular difference in movement direction between two position samples (frames) separated by a 100ms temporal window. The total distance (*d*) traveled between these two frames was also calculated. The process was repeated for the full recording by sliding a 100ms temporal window across all position frames and calculating these values. Angular displacement and distance traveled were converted to Cartesian coordinates to generate x- and y-displacement values in centimeters, which corresponded to lateral and longitudinal displacements for each frame across the full recording.

Two-dimensional displacements were binned (1cm) and convolved with a two-dimensional Gaussian spanning 3cm. For each neuron, the same process was repeated for displacement values that co-occurred with spike times to generate a spike occupancy map as a function of displacement. Self-motion rate maps were constructed by dividing the spike occupancy map for each neuron by the total time in each displacement bin. Bins occupied for less than 267ms were removed from analyses as they typically were observed at extreme displacement values. Self-motion ratemaps for each neuron were additionally constructed independently from interleaved, non-overlapping, 1 second periods throughout the entire session to assess stability of self-motion tuning. For quantification of self-motion tuning relative to a randomized distribution, all aforementioned ratemaps were generated for each neuron 100 times after randomly shifting the spike train relative to position.

Self-motion ratemaps were quantified for their stability, left-versus right-turning preference, and speed modulation. First, stability of self-motion tuning was quantified by correlating self-motion ratemaps generated from non-overlapping periods for each neuron (Figure 2c). Next, turning biases for clockwise versus counterclockwise movements (LvR FR ratio, Figure 2d) were quantified by computing the ratio of summated firing for all displacement bins on the right side and left side of the zero vertical line, respectively. Finally, speed modulation was assessed by finding the mean firing rate as a function of longitudinal displacement (i.e. averaging over columns of self-motion ratemaps) and correlating with the y-displacement value (Figure 2e). All self-motion ratemap quantification was repeated for all randomized self-motion ratemaps as mentioned above. Stability and turn-bias quantification were computed from displacement bins that were occupied in both self-motion ratemaps or both halves of an individual self-motion ratemap. All quantification was completed on non-smoothed self-motion ratemaps.

#### Generalized linear models

In order to more directly test to what degree these neurons represented egocentric compared to allocentric variables, we adopted a generalized linear model framework. The probability of spiking in a given behavioral frame (33Hz) is described by an inhomogeneous Poisson process, where the probability of spiking in a given frame is described by the time varying variable λ:

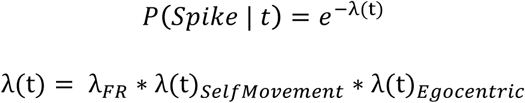

Where:

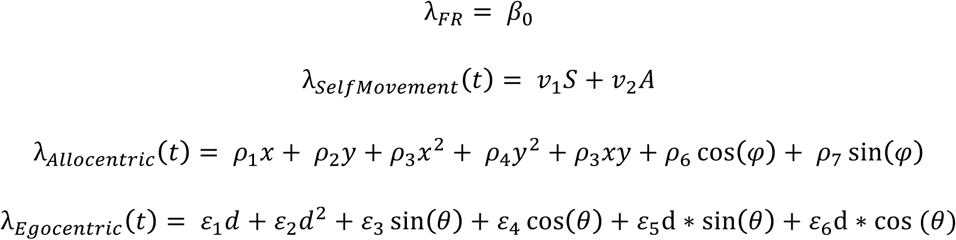

Where *β*_0_ defines the baseline firing rate of the neuron. All subscripted variables are fit coefficients weighting the other (time-varying) variables. S is the running speed of the animal and A is the angular displacement of the animal, as described above. X and y are measurements of the animal’s position in the environment in pixels, and φ is the movement direction. Finally, d is the animal’s distance from the center of the environment, and θ is the egocentric angle to the center of the environment.

Coefficients were determined by fitting to maximize log-likelihood (MATLAB function ‘glmfit’) of the experimental spike train given the behavioral variables. For statistical tests, some number of the coefficients were set to zero, giving a log-likelihood for the reduced model. The difference in likelihood for the full versus reduced model was compared to a Chi-Square distribution (degrees of freedom equal to the number of coefficients set to zero) to generate an analytic p-value. While theoretically the change in log-likelihood should follow a Chi-Square distribution, this is only the case when the spike train has been fit very well (eg: including all neuron-neuron coupling terms). In line with previous approaches, we therefore also compared the change in log-likelihood in two additional manners. First, we compared the change in log-likelihood to that from 1000 randomly shuffled spike train, giving an empirical null-distribution. Secondly, when comparing the relative effects of two variables (that is, comparing two reduced models to each other), we can derive the difference in Akaike Information Criteria (dAIC) for each of the reduced models, and compare their magnitudes. Example spike trains for each model were generated by evaluating lambda for each behavioral time point (‘glmeval’) and using this as the input to a random Poisson Generator (‘poissrnd’).

#### Assessment of theta-phase modulation

For each experimental session an LFP channel was identified that was qualitatively noise free. The LFP signal was filtered in the theta frequency range (6-10Hz) and the phase for each spike from each neuron was estimated as the instantaneous phase angle of the corresponding Hilbert transform (‘hilbert’ in Matlab). For each neuron, the MRL and MRA were calculated on the full spike phase distribution using the circular statistics toolbox (Berens et al., 2009; MRL, ‘circ_r’; MRA, ‘circ_mean’). We next randomly shifted the spike train relative to theta phase 100 times for each neuron to generate a null distribution of MRL values. RSC neurons with MRLs greater than the 95^th^ percentile of the full distribution of randomized MRL values were determined to be theta-phase locked.

#### Assessment of theta rhythmic spiking

Spike train autocorrelograms were estimated by generating a histogram of temporal lags between spikes in a 400ms temporal window discretized into 20ms bins. For each neuron, the power spectrum of the autocorrelogram was computed using the Fourier transform (‘fft’ in Matlab) and the peak in the theta frequency range was identified (if it existed). If the mean power within 1Hz of this theta peak was 50% greater than the mean power for the full power spectrum, the neuron was determined to exhibit intrinsic theta spiking.

#### Gaussian Mixture Models

Preferred orientation and preferred distance estimates were modeled as mixtures of Gaussian distributions using orders from 1 to 10 (‘fitgmist’ in Matlab). Optimal models were identified as those that minimized the Akaike information criterion (AIC). The mean of each Gaussian component is reported and a probability distribution function of the optimal model was generated to visualize mixture model fit (‘pdf’ in Matlab).

#### Statistics

Unless otherwise stated, non-parametric tests with a p-value threshold at 0.05 were utilized for all statistical comparisons. Median and interquartile range are provided for all distributions in which comparisons were made.

#### Data and code availability

The toolbox used for alignment of behavioral and spike data, along with basic analysis is available at https://github.com/hasselmonians/CMBHOME. The toolbox used for egocentric boundary rate map generation and related analyses is available at https://github.com/hasselmonians/rscOpenFieldTuning (to be made public before journal acceptance). The toolbox used for GLM fits, evaluation, and spike-train generation is available at https://github.com/wchapman/pippin (to be made public before journal acceptance).

